# Causal contributions to sensory-based decision-making by cell-type specific circuits in the tail striatum

**DOI:** 10.1101/2022.07.30.502110

**Authors:** Lele Cui, Shunhang Tang, Kai Zhao, Jingwei Pan, Zhaoran Zhang, Bailu Si, Ning-long Xu

## Abstract

The striatum comprises distinct types of neurons giving rise to the direct and indirect basal ganglia pathways and local circuits. A large amount of work has been focusing on cell-type specific striatal circuits in the context of movement control, proposing several models on their functional roles. But it remains to be elucidated how the cell-type specific striatal circuits contribute to decision-making behavior and whether the existing models apply. Here, we investigate the causal roles of the cell-type specific circuits in the posterior tail of the dorsal striatum (TS) of mice in an auditory-guided decision-making behavior. Transient unilateral activation of the direct- or indirect-pathway striatal spiny projection neurons (dSPNs or iSPNs) both biased decisions in opposite directions. These effects, however, were not due to a direct influence on movement, but was specific to the decision period preceding action execution. Optogenetic inactivation of dSPNs and iSPNs revealed their opposing causal contributions to decisions. At the local circuit level, simutaneous optical recording and manipulation of dSPNs and iSPNs revealed their antagnizing interactions. Inactivation of PV interneurons, a common inhibitory input to both dSPNs and iSPNs, facilitated contraversive choices, supporting a causal contribution of coordinated striatal circuits. Using a neural circuit model, we further demonstrated the computational implemenation of the causal circuit mechanism. Our results indicate that while the causal roles of the cell-type specific striatal circuits in decision-making largely agree with classic models in movement control, they show decision task-related specificity involving local circuit coordination.

## Introduction

The striatum in the mammalian brain comprises specific types of projection neurons and local interneurons, which give rise to the two major basal ganglia pathways and form local striatal circuits. The functions of specific types of striatal neurons have been extensively studied in the context of movement control (1–8). In the classic view, the two major types of projection neurons (dSPNs and iSPNs) oppositely regulate movements, as postulated by the classic ‘rate model’, with the dSPNs promoting movements while the iSPNs suppressing movements (4, 5, 9–11). Recent cell-type specific recordings of striatal neuronal activity during movements showed concurrent activation of both dSPNs and iSPNs during movements (3, 7, 12–14), suggesting that despite their opposing roles, the coordination between the two pathways could be important for producing desired movement (2, 5, 15, 16). However, whether and how these postulations apply to more cognitive level behavior, such as decision-making, remain largely unclear. During perceptual decision-making, for instance, different actions need to be selected flexibly based on varying sensory information before motor execution. An important question is how the activity of dSPNs and iSPNs contributes to the sensorimotor decisions beyond action initiation. In addition, the local PV interneurons provide common inhibition to both dSPNs and iSPNs, representing an important local circuit element. While the PV interneurons’ contribution to certain action-based learning has been previously probed (6), how the PV interneurons contribute to decision-making via their inhibitory control of both pathways is still unclear. Understanding the contribution of PV interneurons can be informative of how the two pathways coordinate during decision-making.

Different subregions of the dorsal striatum receives topographically organized cortical and thalamic input (17, 18), as well as the dopaminergic input (19, 20), and may serve distinct functions (16, 21), particularly in different types of cognitive behaviors. For instance, the anterior dorsal striatum (ADS) was shown to be involved in an Poisson-click evidence accumulation task, while dorsal medial striatum (DMS) was implicated in reward-based decision-making task (22, 23). The posterior tail of the dorsal striatum (TS) mainly receives sensory corital input (17, 18) as well as distinct dopamine input signaling stimulus novelty and aversiveness (24, 25), and is considered to be involved in sensory-related decision and learning (24, 26, 27). Here we sought to understand the causal contributions of specific cell types in the TS to perceptual decision-making.

We trained mice to perform a 2-choice auditory decision-making task, and systematically performed temporally confined optogenetic activation and inactivation of specific cell types in the TS, including the dSPNs, iSPNs and PV interneurons during task performance. We found that while dSPNs and iSPNs show opposite contributions to choice behavior, the effects were specific to the decision process but not due to a direct influence on movements. Using simutaneous optical recording and manipulation of dSPNs and iSPNs, we show that the two types of striatal projection neurons exhibit antiganizing interactions in a task-dependent manner, suggesting a within-striatum competition. By inactivating PV interneurons, we disinhibited both dSPNs and iSPNs to produce presumed concurrent activation of the two pathways, and found that this manipulation facilitated contraversive choices, suggesting a coordinated causal contributions of dSPNs and iSPNs. Finally, using a neural circuit model we recapitulated our experimental results, demonstrating the computational plausibility of the causal circuit mechanism.

## Results

### Posterior tail of the striatum causally contributes to perceptual decision-making

Accumulating evidence shows that different subregions of the striatum subserve distinct functions (16, 21). The posterior tail of the striatum (TS) receives topographically organized input, primarily from sensory regions (18, 28). The auditory cortical input to TS was shown to be involved in auditory-decision behavior (27). We thus first tested whether the TS is causally involved in auditory-guided decision-making. We trained head-fixed mice to perform a frequency discrimination psychophysics task (29, 30), where mice were required to classify sound stimuli (pure tones between 7 and 28 kHz) as belonging to low or high frequency categories (**Figure 1A**; **Supplementary Figure 1**). Stimulus categorization was quantified as the probability of choosing the left or right lick port in response to various tone frequencies (**Figure 1B**), allowing for psychophysical measurement of perceptual decisions (**Figure 1C**). To examine the causal role of TS, we injected AAV-CAG-ArchT in the TS of C57/BL6J mice and implanted fiber optics above the injection sites. During task performance, photostimulation (532 nm laser) was delivered to unilateral TS to inactivate neurons on a subset (~15%) of trials randomly interleaved with control trials without photostimulation (**Figure 1D; Supplementary Figure 2**). In a group of mice trained to choose left for the low freuqency category and right for the high frequency category (**Figure 1E**), unilateral inactivation of TS significantly decreased the proportion of choices to the lick port contralateral to the photostimulated hemisphere (**Figure 1E**; two-way repeated-measures ANOVA, F_1,9_ = 21.05, p = 1.3×10^−3^ for photostimulation effect; **p < 0.01, Fisher’s LSD post hoc tests; n= 10 sessions from 5 mice; 1209 photostimulation trials, 3843 control trials).

**Figure 1.**
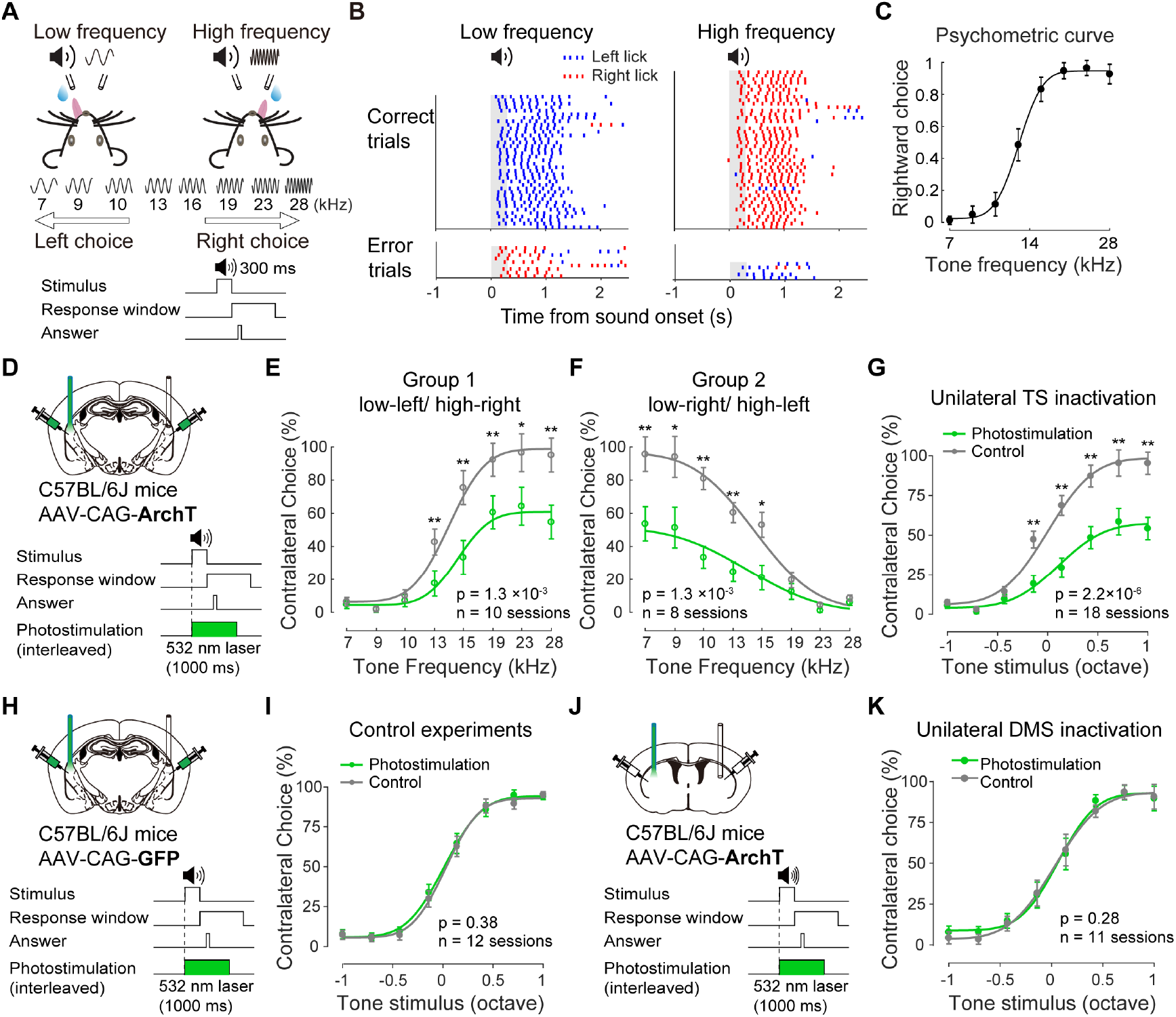
Unilateral inactivation of the posterior tail of the striatum decreased contralateral choices in the auditory-guided decision-making. (A) Top, schematic of the behavioral task. Bottom, temporal structure of the task. (B) Raster plot of lick time from an example session (blue, lick right; red, lick left). Grey shadows indicate the stimulus period. (C) Psychometric function from one example behavioral session. Error bars indicate 95% confidence interval. (D) Schematic showing unilateral photoinactivation of TS using ArchT during auditory decision task. (E) Summarized psychometric function of control trials (grey) and trials with TS inactivation (green), from one groups of mice trained to choose left on low-frequency trials and right on high-frequency trials. n= 10 sessions from 5 mice, F_1,9_ = 21.05, p = 1.3×10^−3^ for photostimulation effect, two-way repeated-measures ANOVA; **p < 0.01, Fisher’s LSD post hoc tests (1209 trials with photostimulation, 3843 control trials). Miss (no lick response) trials were not included. (F) Similar as in (E), but from another group of mice trained to choose right on low-frequency trials and left on high-frequency trials, n= 8 sessions from 4 mice, F_1,7_ = 27.00, p = 1.3×10^−3^ for photostimulation effect, two-way repeated-measures ANOVA; **p < 0.01, *p < 0.05, Fisher’s LSD post hoc tests (989 trials with photostimulation, 3332 control trials). (G) Summary of all sessions shown in (E) and in (F), with percent of contralateral choice (relative to photostimulated hemisphere) as a function of corresponding tone frequencies (in relative octaves). n = 18 sessions from 9 mice, F_1,17_ = 48.75, p = 2.2×10^−6^ for photostimulation effect, twoway repeated-measures ANOVA; **p < 0.01, Fisher’s LSD post hoc tests (2198 trials with photostimulation, 7175 control trials). (H-I) Similar as in (D) and (G), for control group injected with AAV-CAG-GFP in the TS, n = 12 sessions from 3 mice, F_1,11_ = 0.85, p= 0.38 for photostimulation effect, two-way repeated-measures ANOVA (1651 trials with photostimulation, 5329 control trials). (J-K) Similar as in (D) and (G), for optogenetic inactivation of the anterior sub-region of the dorsal striatum, injected with AAV-CAG-ArchT, n = 11 sessions from 6 mice, F_1,10_ = 1.32, p = 0.28 for photostimulation effect, two-way repeated-measures ANOVA (1710 trials with photostimulation, 5697 control trials).

It was previously shown that stimulating the auditory cortical projections in TS biased choices in a sensory-dependent manner (27). In the group of mice shown in **Figure 1E**, the choice direction was correlated with stimulus category, and hence the behavioral effect following photostimulation could be due to an influence on either the contralateral choice or on stimulus categorization of high-frequency sounds. To disambiguate these possibilities, we trained another group of mice with reversed stimulus-choice contingency, with high frequency category corresponding to the left choice and vice versa. We found that unilateral photoinactivation also reduced the contralateral choice, which corresponds to the low frequency category (**Figure 1F**; two-way repeated-measures ANOVA, F_1,7_ = 27.00, p = 1.3×10^−3^ for photostimulation effect; **p < 0.01, *p < 0.05, Fisher’s LSD post hoc tests; n= 8 sessions from 4 mice; 989 photostimulation trials, 3332 control trials), indicating that photoinactivation of TS influenced choice directions independent of stimulus categories (**Figure 1G**; two-way repeated-measures ANOVA, F_1,17_ = 48.75, p = 2.2×10^−6^ for photostimulation effect; **p < 0.01, Fisher’s LSD post hoc tests; n = 18 sessions from 9 mice; 2198 photostimulation trials, 7175 control trials). To control for potential non-specific photo effects, we delivered the same photostimulation in TS of mice expressing GFP (AAV-CAG-GFP in TS), and observed no significant effect on choice behavior (**Figure 1H, I**; two-way repeated-measures ANOVA, F_1,11_ = 0.85, p= 0.38 for photostimulation effect; n = 12 sessions from 3 mice; 1651 photostimulation trials, 5329 control trials).

To assess the specificity of the causal involvement of TS, we also examined the involvement of a more anterior subregion, the dorsomedial striatum (DMS), which is considered to be involved mainly in movement control and reward-based action selection (11, 12, 21–23). We injected AAV-CAG-ArchT in DMS of another group of mice and implanted fiber optics in the same region. Unilateral photoinactivation of DMS during task performance had no significant effects on auditory perceptual decision-making (**Figure 1J, K**; two-way repeated-measures ANOVA; F_1,10_ = 1.32, p = 0.28 for photostimulation effect; n = 11 sessions from 6 mice; 1710 photostimulation trials, 5697 control trials).

Thus, these data support a causal contribution of TS to contralateral choices during auditory perceptual decision-making with a striatal subregion specificity.

### Optogenetic activation of the direct and indirect pathways oppositely regulate perceptual decisions

We next investigate the causal roles of the two major subtypes of striatal projection neurons in the TS, the dSPNs and iSPNs, in auditory decision-making behavior. First, we assessed the effects of optogenetic activation of dSPNs and iSPNs, respectively, on behavioral performance. To activate dSPNs, we expressed Channelrhodopsin-2 (ChR2) in dSPNs by injecting AAV-CAG-FLEX-ChR2 in the TS of D1-Cre mice followed by implanting fiber optics above the injection sites (**Supplementary Figure 3**). During experiments, photostimulation (473 nm laser) was delivered to unilateral TS to activate dSPNs on a subset (~15%) of trials randomly interleaved with control trials (**Figure 2A** and **Supplementary Video 1**; see Methods). We found that activation of dSPNs in TS strongly biased animals’ choices toward the contralateral side relative to the photostimulated hemisphere (**Figure 2B, C**; two-way repeated-measures ANOVA, F_1,21_ = 8.31, p = 8.9×10^−3^ for photostimulation effect; **p < 0.01, Fisher’s LSD post hoc tests; n = 22 sessions from 11 D1-Cre mice; 3083 photostimulation trials, 9491 control trials). This effect was also attributable to an influence on choice direction but not on sensory representation, since photoactivation of dSPNs produced similar contralateral bias in the two separate groups of mice trained with reversed sensory-choice contingencies (**Supplementary Figure 4**).

**Figure 2.**
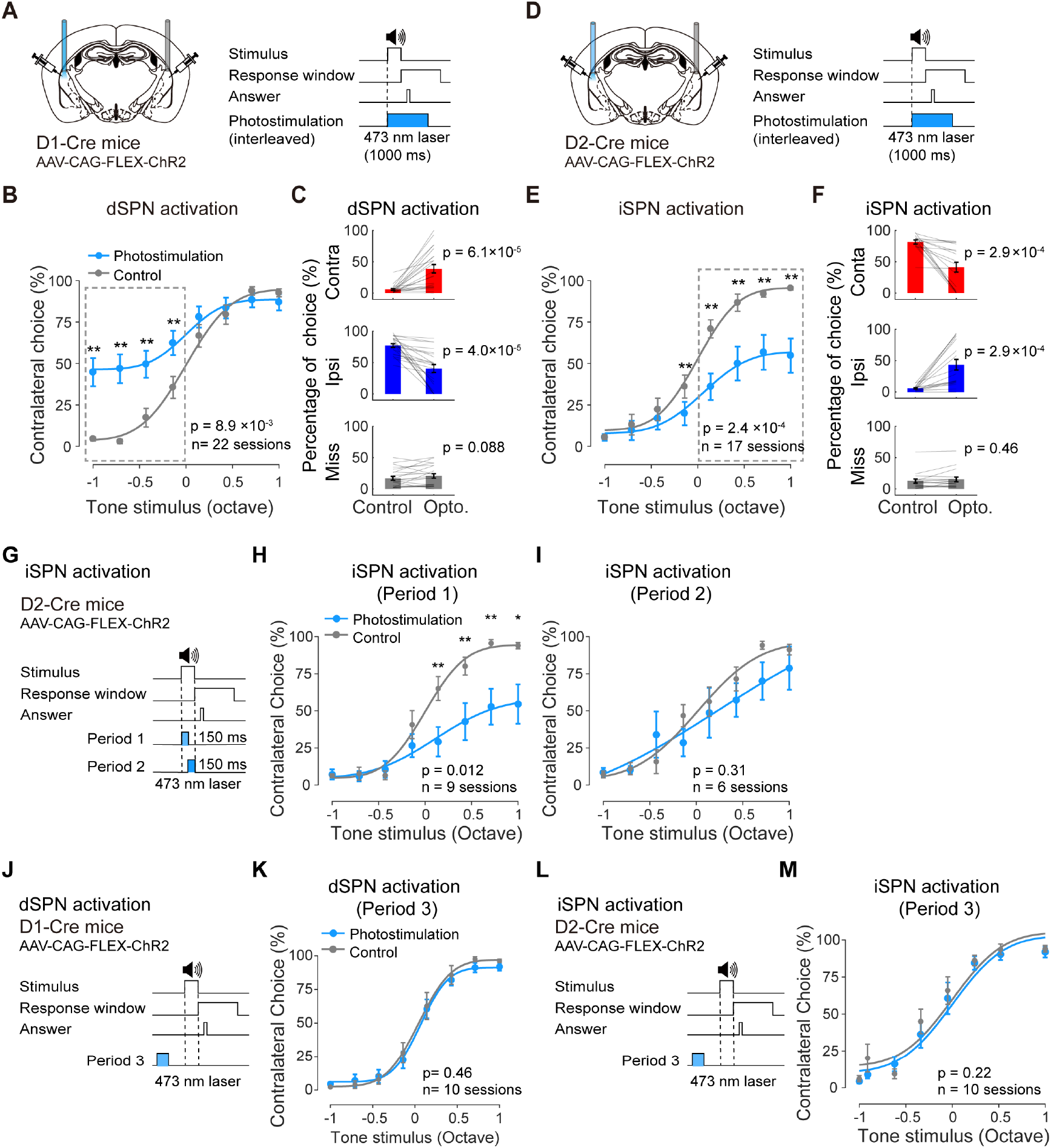
Unilateral optogenetic activation of dSPNs and iSPNs bidirectionally regulate competing choices. (A) Schematic showing experimental configuration for unilateral optogenetic activation of dSPNs. (B) Summarized psychometric functions for control trials (grey) and dSPN activation trials (blue), plotted as the percentage of contralateral choice (relative to photostimulated hemisphere) as a function of tone stimuli in relative octaves. n = 22 sessions from 11 D1-Cre mice; F_1,21_ = 8.31, p = 8.9×10^−3^ for photostimulation effect, two-way repeated-measures ANOVA; **p < 0.01, Fisher’s LSD post hoc tests (3083 trials with photostimulation, 9491 control trials). (C) Summary of choice percentage in trials as indicated in (D) showing contralateral bias, compared between control trials and photostimulation trials for contralateral choice (p = 6.1×10^−5^, from 6±2% to 39±7%, mean±s.e.m.), ipsilateral choice (p = 4.0×10^−5^, from 77±3% to 40±6%), and for miss rate (p = 0.088, from 17±3% to 21±3%), paired two-sided Wilcoxon signed rank test (2114 photostimulation trials, 6214 control trials). (D) Schematic showing unilateral optogenetic activation of iSPNs, similar as in (A). (E) Summarized psychometric functions for control trials and iSPN activation trials, similar as in (B). n = 17 sessions from 9 D2-Cre mice; F_1,16_ = 22.06, p = 2.4×10^−4^ for photostimulation effect, two-way repeated-measures ANOVA; **p < 0.01, Fisher’s LSD post hoc tests (2249 trials with photostimulation, 7666 control trials). (F) Summary of choice percentage in trials as indicated in (E) showing ipsilateral bias, compared between iSPN activation trials and control trials for contralateral choice (p= 2.9×10^−4^, from 82±3% to 41±8%, mean±s.e.m.), ipsilateral choice (p= 2.9×10^−4^, from 6±1% to 43±9%) and for miss rate (p= 0.46, from 13±3% to 15±4%), paired two-sided Wilcoxon signed rank test (1305 trials with photostimulation, 4520 control trials). (G) Schematic showing optogenetic activation of iSPNs at different periods. Blue shades indicate the period of photostimulation. Period 1, a 150 ms window following sound onset; Period 2, a 150 ms window before the end of tone stimulus. (H) Summarized psychometric functions similar as in (E) for iSPN activation in period 1. n = 9 sessions from 6 D2-Cre mice; F_1,8_ = 10.50, p = 0.012 for photostimulation effect, two-way repeated-measures ANOVA; *p < 0.05, **p < 0.01, Fisher’s LSD post hoc tests (1248 trials with photostimulation, 4236 control trials). (I) Summarized psychometric functions similar as in (E) for iSPN activation in period 2. n = 6 sessions from 3 D2-Cre mice; F_1,5_ = 1.27, p = 0.31 for photostimulation effect, two-way repeated-measures ANOVA (732 trials with photostimulation, 2377 control trials). (J) Schematic showing optogenetic activation of dSPNs in period 3 (before sound onset during inter-trial-interval). (K) Similar as in (B) for dSPN activation in period 3. n = 10 sessions from 5 D1-Cre mice; F_1,9_ = 0.60, p =0.46 for photostimulation effect, two-way repeated-measures ANOVA (1538 trials with photostimulation, 3920 control trials). (L) Schematic showing optogenetic activation of iSPNs in period 3 as in (J). (M) Similar as in (E) for iSPN activation in period 3. n = 10 sessions from 3 D2-Cre mice; F_1,9_ = 1.73, p = 0.22 for photostimulation effect, two-way repeated-measures ANOVA (1355 trials with photostimulation, 4281 control trials).

Although it was proposed that the dSPN activation could produce positive reinforcement and could change action values (23, 31), the effects on choice behavior here was unlikely due to a persistent change in motivation level since we delivered photonstimulation trial-by-trial in only ~15% of trials randomly interleaved with control trials, precluding potential contributions from accumulated reinforcement. Indeed, the increase of contralateral choices was not accompanied by any significant change in miss rate (**Figure 2C** and **Supplementary Figure 5B**), suggesting that there were no significant changes in motivation level. The effects on choices were neither due to changes in direct motor control of licking, since the lick rate in individual trials was not significantly different between photostimulation and control trials (**Supplementary Figure 5D**). Thus, dSPN activation specifically promotes the decision-related contralateral choices.

To activate iSPNs we injected AAV-CAG-FLEX-ChR2 in the TS of D2-Cre mice (**Figure 2D**; **Supplementary Figure 3**; see Methods). We found that unilateral activation of iSPNs strongly biased animals’ choices toward the ipsilateral side relative to the photostimulated hemisphere (**Figure 2E, F**; two-way repeated-measures ANOVA, F_1,16_ = 22.06, p = 2.4×10^−4^ for photostimulation effect; **p < 0.01, Fisher’s LSD post hoc tests; n = 17 sessions from 9 D2-Cre mice; 2249 trials with photostimulation, 7666 control trials), an effect opposite to that of dSPN activation. Similarly, we found no significant changes in miss rate during photostimulation (**Supplementary Figure 5C**), ruling out potential influence of motivational state following iSPN activation. Earlier models emphasize the suppressive effect on movement by the indirect pathway (5, 32). Here we found that while activation of iSPNs reduced contralateral choices, it also increased ipsilateral choices (**Figure 2F**). Together with the unchanged miss rate, this suggests that the effect of iSPN activation was not simply a general suppression of contralateral actions, but was part of the competition process for selecting actions during decision-making.

While dSPN activation did not affect lick rate (**Supplementary Figure 5D**), activation of iSPN was accompanied by a slightly but significant decrease in the contralateral lick rate following sensory stimulus (**Supplementary Figure 5E**). We asked whether this effect represents a decision-specific change in the contralateral choice or reflects a direct influence on lick movement. We thus applied more precisely confined photoactivation to iSPNs during task performance. Since we observed that mice often make correct answer lick before the end of sound stimulus (~150 ms after sound onset, **Supplementary Figure 6A**), decisions could be formed well before the end of sound stimulus. We reasoned that restricting the photoactivation of iSPNs within the first half of the stimulus duration (a 150 ms window after sound onset) would mainly affect decision formation before motor execution, avoiding the majority of answer licks (**Figure 2G**, period 1). We found that this photoactivation strongly biased animals’ choice toward the ipsilateral side (**Figure 2H**; two-way repeated-measures ANOVA, F_1,8_ = 10.50, p = 0.012 for photostimulation effect; *p < 0.05, **p < 0.01, Fisher’s LSD post hoc tests; n = 9 sessions from 6 D2-Cre mice; 1248 photostimulation trials, 4236 control trials), similar to the effect found for the photoactivation covering both stimulus and response epochs (**Figure 2E**). In contrast, when we restricted photoactivation to the second half (duration, 150 ms) of the sound stimulus period where mice often made putative post-decision answer licks (**Figure 2G**, period 2), the choice behavior was largely unaffected (**Figure 2I**; two-way repeated-measures ANOVA, F_1,5_ = 1.27, p = 0.31 for photostimulation effect; n = 6 sessions from 3 D2-Cre mice; 732 trials with photostimulation, 2377 control trials). Thus, the effect on choice behavior following iSPN activation was due to specific influence on perceptual decisions but not due to direct influence on lick movement.

To further rule out any potential effects of photoactivation before sensory stimulus, we delivered the photoactivation in an inter-trial-interval epoch before the sound stimulus onset for both dSPNs (**Figure 2J**) and iSPNs (**Figure 2L**). We found no significant changes in the behavioral performance for dSPN activaiton (**Figure 2K**; two-way repeated-measures ANOVA; F_1,9_ = 0.60, p =0.46 for photostimulation effect; n = 10 sessions from 5 D1-Cre mice; 1538 trials with photostimulation, 3920 control trials) and for iSPN activation (**Figure 2M**; two-way repeated-measures ANOVA, F_1,9_ = 1.73, p = 0.22 for photostimulation effect; n = 10 sessions from 3 D2-Cre mice; 1355 photostimulation trials, 4281 control trials).

Taken together, these results indicate that both dSPNs and iSPNs in TS oppositely regulate sensory-based decision-making with a high degree of specificity. While the classic models predict general opposing roles of dSPNs and iSPNs in movement control, here we provide evidence extending such predictions to perceptual decision-making in the striatal subregion concordant with its input topography.

### Optogenetic inactivation reveals opposing contributions to choice behavior by dSPNs and iSPNs

While optogenetic activation often produce strong behavioral effects, to reveal the contributions by endogenous activity of different types of striatal neurons would require inactivation of specific cell types during task performance. We thus further use optogenetic inactivation to examine the involvement of dSPNs and iSPNs, respectively, in sensory-based decision-making. To inactivate dSPNs, we injected AAV-Ef1α-DIO-eNpHR3.0 in the TS of D1-Cre mice to express eNpHR3.0 in dSPNs (**Supplementary Figure 7**) and delivered unilateral photostimulation (593 nm) covering the sensory and decision epochs on interleaved trials (**Figure 3A**). We found that inactivation of dSPNs significantly biased animals’ choice toward the ipsilateral side relative to the photostimulated hemisphere (**Figure 3B, C**; two-way repeated-measures ANOVA, F_1,1_1 = 13.39, p = 3.8×10^−3^ for photostimulation effect; *p < 0.05; **p < 0.01, Fisher’s LSD post hoc tests; n = 12 sessions from 6 D1-Cre mice; 1594 trials with photostimulation, 5271 control trials). To inactivate iSPNs, we injected AAV-Ef1α-DIO-eNpHR3.0 in the TS of D2-cre mice to express eNpHR3.0 in iSPNs (**Figure 3D**). We found that iSPN inactivation led to a slight but significant bias of animals’ choice toward the contralateral side (**Figure 3E, F**; two-way repeated-measures ANOVA, F_1,19_ = 21.85, p = 1.7×10^−4^ for photostimulation effect; *p < 0.05, Fisher’s LSD post hoc tests; n = 20 sessions from 10 D2-Cre mice; 2998 photostimulation trials, 8303 control trials). To understand the relatively weak behavioral effects, especially following iSPN inactivation, we expressed the soma-targeted *Guillardia theta* anion-conducting channelrhodopsin-2 (stGtACR2), a more potent activity suppressor (33), in iSPNs, and delivered blue laser (473 nm) to inactivate iSPNs during task performance (**Figure 3G**). We found a significant contralateral biasing effect following iSPN inactivation by stGtACR2 (**Figure 3H, I**; two-way repeated-measures ANOVA, F_1,15_ = 11.72, p = 3.8×10^−3^, for photostimulation effect; *p < 0.05, **p < 0.01, Fisher’s LSD post hoc tests; n = 16 sessions from 8 D2-Cre mice; 6318 photostimulation trials, 9334 control trials), an effect similar to that following inactivation using eNpHR3.0. Unilateral inactivation of dSPNs or iSPNs did not significantly influence miss rate, precluding potential effects on motivation (**Figure 3C, F** and **I; Supplementary Figure 8A-C**). The lick rate during task performance was largely unchanged following inactivation of dSPNs or iSPNs, ruling out a direct influence on lick movement (**Supplementary Figure 8D-F**). Thus, unilateral inactivation of dSPNs or iSPNs revealed opposing contributions to sensory-based decision-making by the endogenous activity of these cell types.

**Figure 3.**
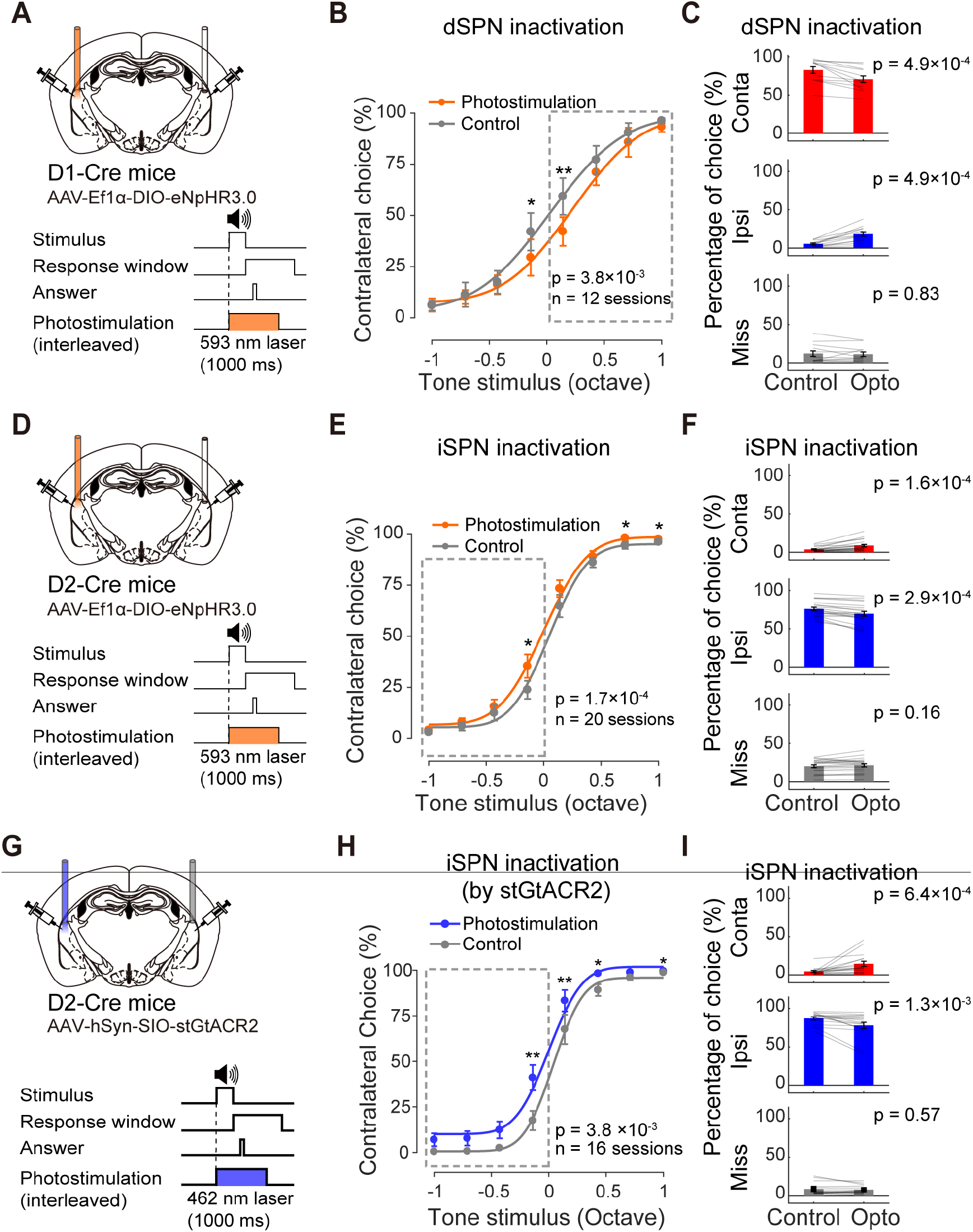
Unilateral optogenetic inactivation of dSPNs and iSPNs supports their complementary contributions to behavioral choices. (A) Schematic showing experimental configuration for unilateral optogenetic inactivation of dSPNs using eNpHR3.0. (B) Summarized psychometric functions for unilateral dSPN inactivation. n = 12 sessions from 6 D1-Cre mice; F_1,11_ = 13.39, p = 3.8×10^−3^ for photostimulation effect, two-way repeated-measures ANOVA; *p < 0.05; **p < 0.01, Fisher’s LSD post hoc tests (1594 trials with photostimulation, 5271 control trials). (C) Summary of choice percentage in trials as indicated in (B) to assess ipsilateral bias. Choices are compared between control trials and photostimulation trials for contralateral choice (p = 4.9×10^−4^, from 82±4% to 70±4%, mean±s.e.m.), ipsilateral choice (p = 4.9×10^−4^, from 5±1% to 18±3%), and for miss rate (p = 0.83, from 12±4% to 11±3%), paired two-sided Wilcoxon signed rank test. (D) Schematic for unilateral iSPN inactivation using eNpHR3.0. (E) Summarized psychometric functions for unilateral iSPN inactivation. n = 20 sessions from 10 D2-Cre mice; F_1,19_ = 21.85, p = 1.7×10^−4^ for photostimulation effect, two-way repeated-measures ANOVA; *p < 0.05, Fisher’s LSD post hoc tests (2998 trials with photostimulation, 8303 control trials). (F) Summary of choice percentage in trials as indicated in (E) to assess contralateral bias, compared between iSPN inactivation trials and control trials for contralateral choice (p = 1.6×10^−4^, from 4±1% to 9±2%, mean±s.e.m.), ipsilateral choice (p = 2.9×10^−4^, from 76±2% to 70±3%) and miss (p = 0.16, from 21±2% to 22±2%), paired two-sided Wilcoxon signed rank test. (G) Schematic for unilateral iSPN inactivation using stGtACR2. (H) Summarized psychometric functions for unilateral iSPN inactivation using stGtACR2. n = 16 sessions from 8 D2-Cre mice; F_1,15_ = 11.72, p = 3.8×10^−3^, for photostimulation effect, two-way repeated-measures ANOVA; *p < 0.05, **p < 0.01, Fisher’s LSD post hoc tests (6318 trials with photostimulation, 9334 control trials). (I) Summary of choice percentage in trials as indicated in (H) to assess contralateral bias, compared between iSPN inactivation and control trials for contralateral choice (p = 6.4×10^−4^, from 5±1% to 15±3%, mean±s.e.m.), ipsilateral choice (p = 1.3×10^−3^, from 87±2% to 78±4%), and for miss rate (p = 0.57, from 8±2% to 8±1%; paired two-sided Wilcoxon signed rank test.

### Concurrent disinhibition of dSPNs and iSPNs promotes contralateral choice

The local PV interneurons in the striatum provide common inhibition to both dSPNs and iSPNs, representing an important circuit element coordinating the two pathways. While the function of striatal PV interneurons in certain action-based learning has been previously probed (Owen et al., 2018), how the PV interneurons contribute to decision-making via their inhibitory control of both pathways has not been demonstrated yet. We thus performed optogenetic inactivation of the PV interneurons in TS by expressing eNpHR3.0 in these neurons using AAV-Ef1a-DIO-eNpHR3.0 in PV-Cre mice (**Figure 4A**; **Supplementary Figure 9A**). We found that unilateral inactivation of PV interneurons in the TS significantly biased choices toward the contralateral side (**Figure 4B,C**; two-way repeated-measures ANOVA; F_1,13_ = 13.09, p = 3.1×10^−3^ for photostimulation effect; **p < 0.01, Fisher’s LSD post hoc tests; n = 16 sessions from 8 PV-Cre mice; 1548 photostimulation trials, 4248 control trials), without any significant effects on miss rate and lick rate (**Supplementary Figure 10**), indicating that PV interneurons are causally involved in decision-making behavior, suppressing contralateral choices or promoting ipsilateral choices.

**Figure 4.**
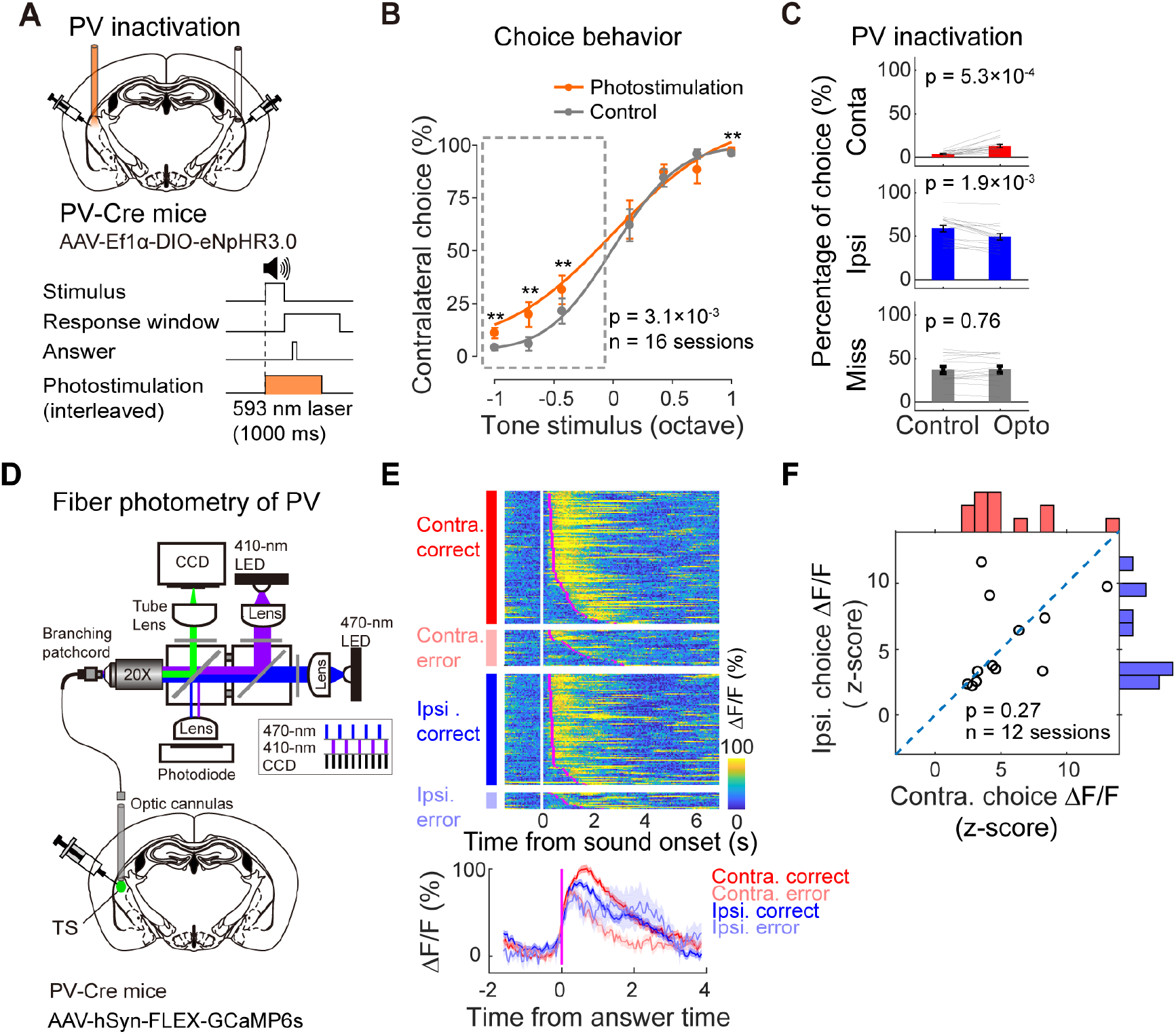
Inactivation of PV interneurons in the TS biased choices towards the contralateral side. (A) Schematic showing unilateral optogenetic inactivation of PV interneuron in TS. (B) Summarized psychometric functions following unilateral PV interneuron inactivation. n = 16 sessions from 8 PV-Cre mice; F_1,13_ = 13.09, p = 3.1×10^−3^ for photostimulation effect, two-way repeated-measures ANOVA; **p < 0.01, Fisher’s LSD post hoc tests (1548 trials with photostimulation, 4248 control trials). (C) Summary of choice percentage in trials as indicated in (B), compared between PV inactivation trials and control trials for contralateral choice (p = 5.3×10^−4^, from 4±1% to 13±2%, mean±s.e.m.); ipsilateral choice (p = 1.9×10^−3^, from 59±4% to 49±4%); miss (p = 0.76, from 37±4% to 37±4%), paired two-sided Wilcoxon signed rank test. (D) Schematic showing fiber photometry recording from PV interneurons in TS. (E) Calcium signals of PV interneurons in the TS from an example behavior session. Upper, color raster plot of calcium signals from individual trials sorted according to the four trial types; lower, mean traces averaged for different trial types. (F) Summarized responses of PV interneurons compared between trials with contralateral and ipsilateral choices. n = 12 sessions from 6 mice, p = 0.27, paired two-sided Wilcoxon signed rank test.

A simple explanation for such causal effect is that PV interneurons take actions by carrying choice information, which would predict that they show stronger responses upon ipsilateral choice but weaker responses upon contralateral choices. We thus examined the activity of striatal PV interneurons during task performance using fiber photometry to record calcium signals in a cell-type specific manner. We expressed GCaMP6s in PV interneurons by injecting AAV-hSyn-FLEX-GcaMP6s in the TS of PV-Cre mice, and implanted fiber optics above the injection sites (**Figure 4D** and **Supplementary Figure 9B**). We found that PV interneurons in TS are strongly activated during the choice period (**Figure 4E**), suggesting an active participation in the operation of striatal circuits during choice behavior. However, contrary to the simple prediction, we found that PV interneurons showed comparable responses to contra- and ipsilateral choices without significant choice preferences (**Figure 4F**; paired two-sided Wilcoxon signed rank test, p = 0.27, n = 12 sessions from 6 mice). This implies that rather than directly encoding choice information, PV interneurons could contribute to choice behavior by regulating the balance and competition between dSPNs and iSPNs in the local circuitry, and the choice bias following PV inactivation likely resulted from the concurrently elevated activity in both dSPNs and iSPNs.

### Antagonizing effect exerted by iSPNs on dSPNs during decision-making behavior

The effect of PV interneurons on choice behavior is likely via its inhibition on other striatal cell types, implying that the interaction between different cell types can be an important factor in the striatal circuit operation. While the dSPNs and iSPNs exert their effects mainly through their projections to downstream regions, previous studies revealed antiganizing interactions between these two cell types in *ex vivo* preparation (34, 35). We thus probed the interaction between dSPNs and iSPNs during decision-making behavior using simutaneous optical recording and manipulations (**Supplementary Figure 11**). To achieve cell-type specificity in both manipulation and recording simulaneously, we exploited both genetic-based and projection-based cell-type targeting approaches. To activate iSPNs, we injected AAV-hSyn-FLEX-ChrimsonR in TS of D2-cre mice to express ChrimsonR in iSPNs and implanted fiber optics in the same region. To simultaneously record activity from dSPNs in the same animals, we took advantage of the target specificity of striatal projection neurons, in that only dSPNs but not iSPNs project to the substantia nigra pars reticulata (SNr). We injected AAV-hSyn-GCaMP6s in TS to express GCaMP6s non-selectively in TS, and implanted a fiber optics in the ipsilateral SNr to record calcium signals only from axons originating from dSPNs in TS (**Figure 5A; Supplementary Figure 11A**). Consistent with previous fiber photometry recording from the somatic region of dSPNs (36), we found the axonal activity of dSPNs show higher responses in contralateral trials than ipsilateral trials (**Supplementary Figure 12**). Interestingly, photoactivation of iSPNs resulted in a significant reduction in dSPNs axonal responses in contralateral trials, but not in ipsilateral trials (**Figure 5C**; paired two-sided Wilcoxon signed rank test, contralateral trials, %Δ*F/F* from 2.99±1.01 to 1.80±0.36, *p* = 0.024; ipsilateral trials, %Δ*F/F* from 1.68±0.31 to 2.06±0.34, p= 0.15; n = 11 sessions from 4 D2-Cre mice), suggesting a behavioral contex-dependent interaction the two cell types.

**Figure 5.**
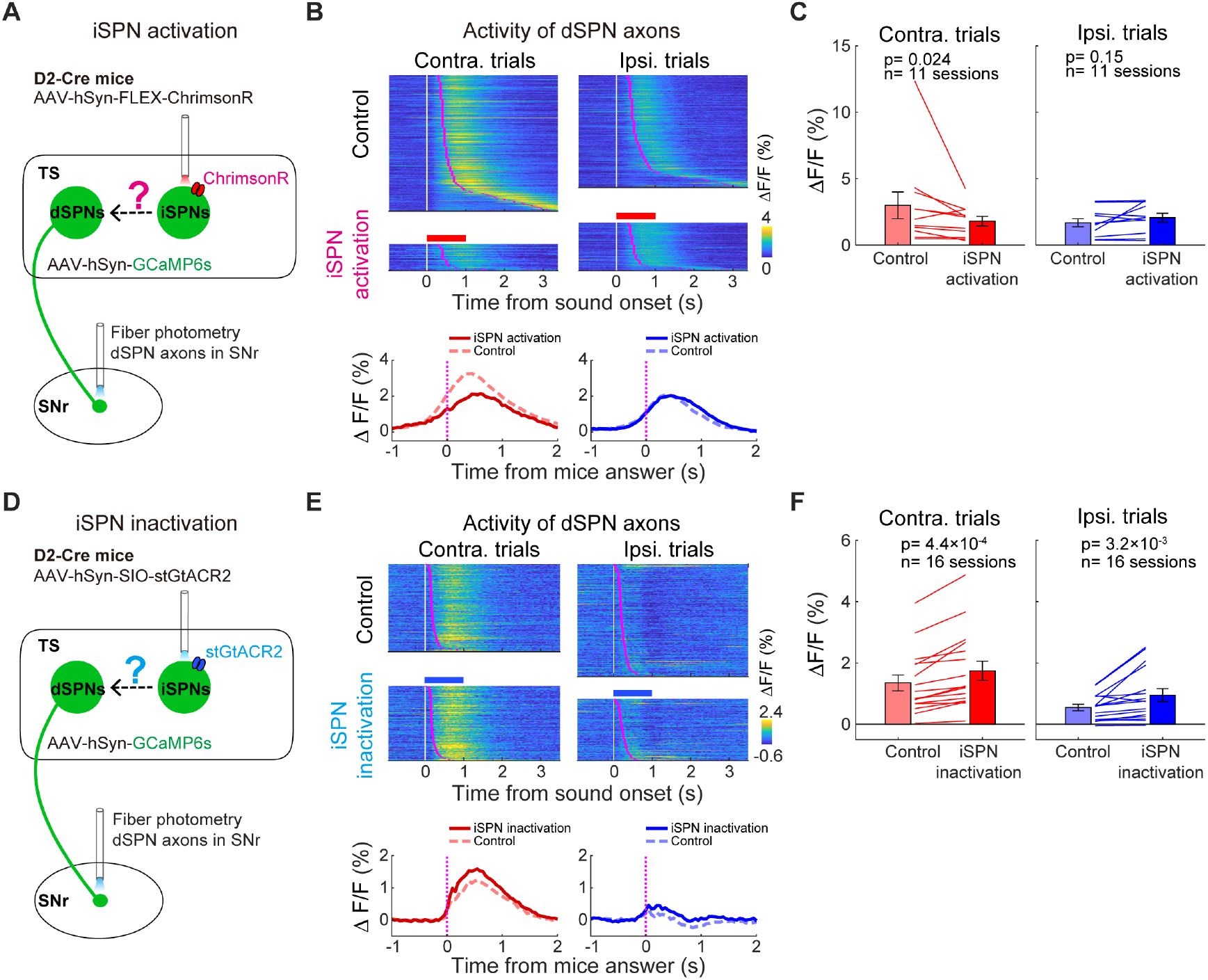
Antagonizing effect exerted by iSPNs on dSPNs during auditory-guided decision-making. (A) Experimental configuration for simultaneous fiber photometry recording from dSPN axons in SNr and optogenetic activation of iSPN in TS. (B) Calcium signals from dSPN axons in SNr in an example behavioral session. Trials with photostimulation and control trials were separated (only correct trials were included). Upper, color raster plot; lower, mean traces averaged for different trial types. (C) Summary of dSPN axonal activity compared between control trials and photostimulation trials as in (B). Contralateral trials, Δ*F/F*% from 2.99±1.01 to 1.80±0.36, mean±s.e.m, p=0.024; ipsilateral trials, Δ*F/F*% from 1.68±0.31 to 2.06±0.34, p=0.15. Each line represents one recording session. n = 11 sessions from 4 D2-Cre mice, paired two-sided Wilcoxon signed rank test. (D) Experimental configuration for simultaneous fiber photometry recording from dSPN axons in SNr and iSPN optogenetic inactivation in TS using stGtACR2. (E) Calcium signals from dSPN axons in SNr with or without iSPN inactivation from an example behavioral session, similar as in (B). (F) Summary of dSPN axonal activity compared between trials with and without photostimulation as in (E). Contralateral trials, Δ*F/F*% from 1.35±0.26 to 1.74±0.32, mean±s.e.m, p = 4.4×10^−4^; ipsilateral trials, from Δ*F/F*% 0.55±0.11 to 0.95±0.21, p = 3.2×10^−3^. n = 16 sessions from 8 D2-Cre mice, paired two-sided Wilcoxon signed rank test.

We used the similar approach to inactivate iSPNs while recording activity in dSPN axons (**Figure 5D**). We co-injected AAV-hSyn-SIO-stGtACR2 and AAV-hSyn-GCaMP6s in TS of D2-Cre mice to express stGtACR2 only in iSPNs and GCaMP6s non-selectively. We implanted fiber optics in TS for photostimulation and in SNr for photometry. We found that optogenetic inactivation of iSPNs increased the response level of dSPN axons (**Figure 5E,F**; paired two-sided Wilcoxon signed rank test, contralateral trials, %Δ*F/F* from 1.35±0.26 to 1.74±0.32, p=4.4×10^−4^; ipsilateral trials, from 0.55±0.11 to 0.95±0.21, p=3.2×10^−3^; n = 16 sessions from 8 D2-Cre mice), suggesting that iSPNs normally exert an inhibitory effect on dSPN activity. Thus, iSPNs exert an antagonizing effect on dSPN activity, suggesting a within-striatum competition mechanism between different subtypes of projection neurons during decision-making behavior.

### Computational implementation of the coordination of striatal cell-type specific circuits

Our experimental data reveal the causal roles of cell type-specific circuits in TS in sensory-based decision-making. To further understand how the causal effects of different cell types coordinate at circuit level, we constructed a computational model of the TS circuits with different cell types (**Figure 6A**; **Supplementary Table 1**). In this model, the sensory decisions are derived from cortex and sent to dSPNs, iSPNs and striatal PV neurons via cortico-striatal projections (37). Within the local striatal region, PV neurons provide inhibitory input to both dSPNs and iSPNs, and the two subtypes of SPNs provide mutual inhibitory input to each other (34, 35). The dSPN and iSPN populations provide either inhibitory or excitatory (indirectly) input to ipsilateral SNr through the direct and indirect pathways respectively (9, 38). The left and right SNr populations compete with each other, and the dominant SNr population determines the left or right choices. The response preferences and amplitude of dSPN and iSPNs were derived from the results in our separate study using cell-type specific two-photon imaging from TS under the same behavior task (39), which shows that both dSPNs and iSPNs contain divergent subpopulations preferring either ipsi- or contralateral choices, with higher response strength in contralateral preferring neurons, and the contralateral dominance was stronger in dSPNs than in iSPNs (**Figure 6B**).

**Figure 6.**
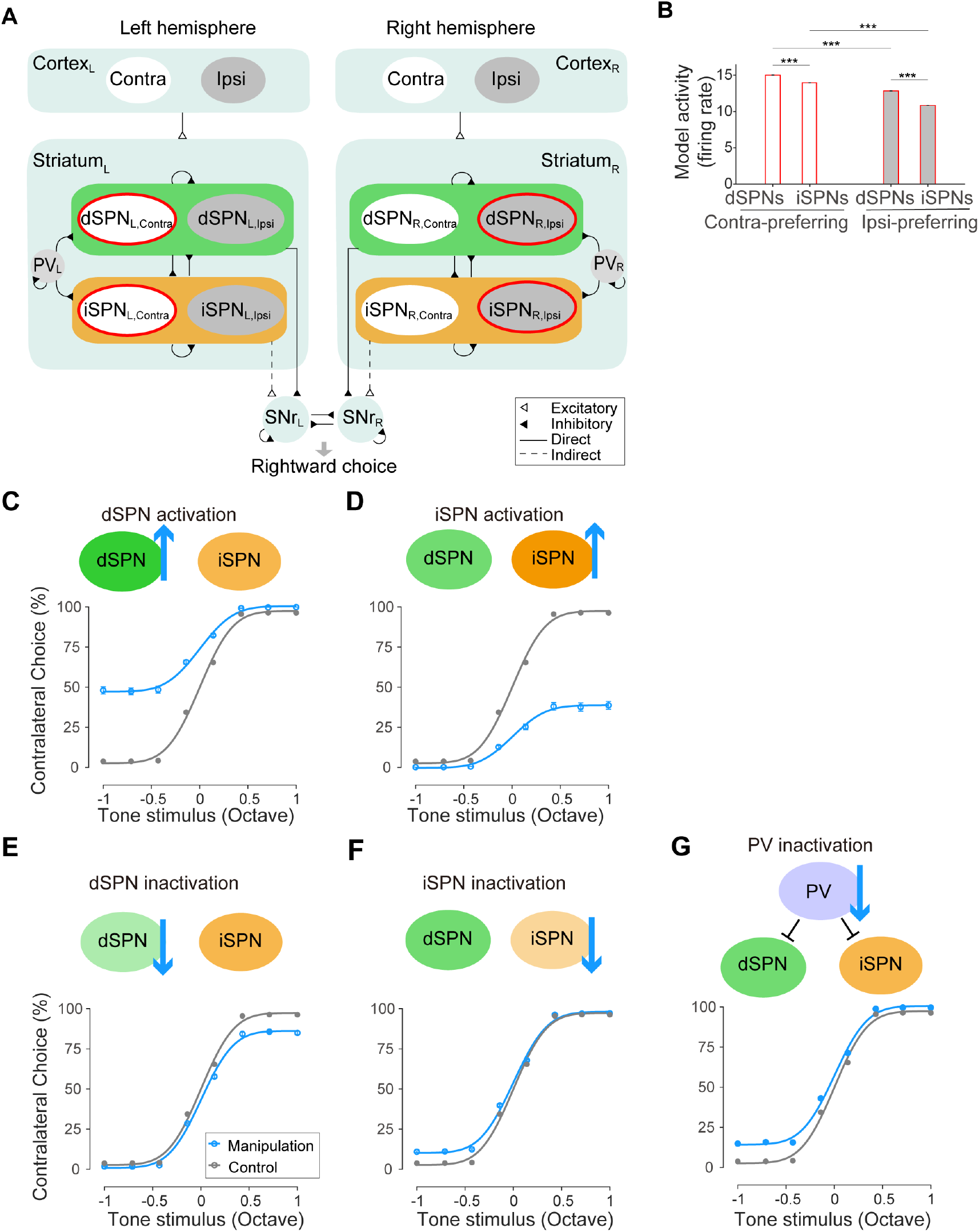
A neural network model recapitulates the causal roles of cell-type specific striatal circuits in perceptual decision behavior. (A) Schematic showing the network model organization. L, left hemisphere; R, right hemisphere; Contra, contralateral preferring neuronal population; Ipsi, ipsilateral preferring neuronal population. See Methods. (B) Responses of modeled dSPNs and iSPNs with contralateral or ipsilateral preference in preferred trial types. ***, p <0.001, two-sided Wilcoxon rank sum test. (C-G) Model simulation of behavioral effects of optogenetic manipulations. Model outputs of choice behavior from control trials were shown in grey, and those from trials with unilateral dSPN activation (C), iSPN activation (D), dSPN inactivation (E), iSPN inactivation (F) and PV interneuron inactivation (G) were shown in blue. Each curve was obtained from 22 different values (Supplementary table 2).

We asked whether our model could recapitulate the perceptual decision behavior under both control condition and cell-type specific manipulation conditions. First, our model recapitulated the choice behavior with similar psychometric functions as in behavioral data (gray lines in **Figure 6C-G**). Next, we increased or decreased the activity level of different striatal cell types in our model to simulate optogenetic manipulations (**Supplementary Table 2**). We found that unilateral increasing of dSPN activity biased the choice towards the contralateral side, whereas unilateral increasing of iSPN activity biased the choice towards the ipsilateral side (**Figure 6C,D**), which qualitatively captured our experimental results of optogenetic activation (**Figure 2A-F**). Unilateral decreasing of dSPN activity in the model biased the choice towards ipsilateral side, whereas unilateral decreasing of iSPN activity in the model biased the choice towards contralateral side (**Figure 6E-F**), which also captured our optogenetic inactivation results (**Figure 3**). Finally, we simulated the concurrent disinhibition by decreasing the activity of unilateral PV interneurons in the model (**Supplementary Table 2**), which biased the choice towards the contralateral side (**Figure 6G**). This result also captures our experimental observation following PV neuron inactivation (**Figure 4A-C**). Together, our simulation results using the circuit-level model support the computational plausibility of the causal contributions by cell type-specific striatal circuits to sensory-guided decision-making.

## Discussion

The striatal circuitry has been extensively studies in the context of movement control, revealing the respective roles of different types of projection neurons and local interneurons (2, 4–6, 11). This can stem from the long history of neurological research focusing on the prominent association between dysfunctions of the striatal circuitry and motor-related disorders, including Parkinson’s and Huntington’s diseases. However, it has been increasingly recognized that the striatal circuitry also plays key roles in more cognitive level functions and dysfunctions, including decision-making (16, 23, 27, 40–43), and perceptual disturbances associated with schizophrenia (44–47). An important question therefore is how the cell type-specific circuits in striatum contribute to decision-making and perceptual functions, and whether previous models formulated by studying motor functions apply. Here we investigate the causal roles of specific cell types in the tail of dorsal striatum in an auditory decision-making task. For the projecting cell types, dSPNs and iSPNs, optogenetic manipulations revealed their opposing effects on choice behavior with the temporal specificity in the decision-relevant period (**Figure 2** and **Figure 3**). These manipulation results extend the classic notion of the differential roles of the direct and indirect striatal pathways in movement control to the domain of perceptual decision-making with trial-by-trial specificity and sub-trial precision. For local circuits, we target an important local inhibitory neuron, the PV interneurons. While the responses of PV interneurons show balanced selectivity to both contra- and ipsilateral choices, PV neurons exert a significant suppressive effect on the contralateral choice (**Figure 4**), likely reflecting an effect of coordinated co-activation of the two projection cell types. In addition to the common inhibition from PV neurons to dSPNs and iSPNs, we also found that iSPNs exert an antagonizing effect on dSPN activity during task performance (**Figure 5**). Finally, we used a neural circuit model to demonstrate the circuit-level computational plausibility of the experimentally observed causal mechanisms (**Figure 6**).

In the past, various models have been proposed to account for the role of the striatal pathways in movement control (2, 5, 9, 15, 16, 48). The classic rate model postulates opposing role of the dSPNs and iSPNs in movement control, primarily concerning motor initiation. While this model did not provide specific predictions on decision-related action selection, our optogenetic manipulation results provide new evidence for the opposing causal roles of dSPN and iSPN in the perceptual decision process (**Figure 2** and **Figure 3**). In addition, our temporally restricted manipulations highlighted decision-specific roles of these projection neurons in TS (**Figure 2H**), independent of general movement control including drinking related licking behavior (**Supplementary Figure 5**). Recently observed concurrent activation of the direct and indirect striatal pathways during movement (7, 12, 14) suggests that despite their opposing roles in controlling movement, these two pathways are likely to act together via competition or coordination during movement (2, 5, 16), which has been be formulated as “competitive” model (15), or “complementary” model (9, 11, 15, 48). In our PV neuron inactivation experiments, concurrent disinhibition of both dSPNs and iSPNs likely elevated the activity level of both pathways by their endogenous excitatory inputs, which resulted in biased decisions towards the contralateral choices (**Figure 4**). Such behavioral effect is likely to result from the competition between the two pathways that were concurrently active. In accordance with our experimental results, our circuit model shows that with stronger contralateral preference in dSPNs than in iSPNs, suppression of PV interneurons with comparable inhibitory weights on dSPNs and iSPNs, indeed biased choices towards the contralateral side (**Figure 6**). Thus, our results extend the previous models regarding the cell type specific striatal circuits concerning movement control to perceptual decision-making domain, and reveal additional specificity in decision process.

The causal roles of the cell-type specific striatal circuits have been widely studied in various forms of motor function, including locomotion (4, 7, 11), sequential actions (49), and movement kinematics (8). For sensory-guided decision-making, however, the causal roles of specific striatal cell types are less understood. We found that activation of dSPNs and iSPNs elicited strong but oppositely lateralized decision-bias (**Figure 2**), corroborating the general notion of the functional opponency of the direct and indirect striatal pathways. By precisely restricting photostimulation in different sub-trial time windows, we demonstrate that striatal projection neurons in TS play specific causal roles in sensory-guided decision process rather than in ensuing movement (**Figure 2**). In addition to activation (23, 43), it is important to also examine the effect of inactivation to reveal the causal contributions of endogenous neuronal activity. Here, we found that optogenetic inactivation of dSPNs and iSPNs led to opposite lateralized biasing of choice behavior with reversed directions than the activation effects (**Figure 3**), revealing the causal involvement of the two striatal pathways in decision-making behavior. While being an important circuit element, the causal contribution of striatal PV interneurons to behavior is even less understood. Whereas previous disruption of striatal PV neurons appeared not to directly affect motor performance (6), here using optogenetic inactivation of PV interneurons in TS, we unraveled a clear biasing effect on choice behavior (**Figure 4**), reflecting its important role in regulating the coordination of striatal circuits.

While another type of local interneurons in the striatum, the cholinergic interneurons (CINs), has been considered to also coordinate the local striatal circuits, they were reported to mainly exert effects in learning (16, 50), and recently in locomotion (51). Its role in decision-making, particularly in TS, could be complex and can be a subject for future studies. In addition, while the dSPNs and iSPNs in TS receive excitatory input primarily from sensory cortex and thalamus, they are also under heavy influence of midbrain dopaminergic input (19). The interactions between the excitatory inputs and dopaminergic input in TS may play critical roles in regulating perceptual and cognitive functions (20, 24, 45–47). The underlying circuit and plasticity mechanisms for such interactions in TS during decision-making and learning are also important subjects for future investigation.

## Materials and Methods

### Subjects

All experimental aspects were carried out in compliance with the procedures approved by the Animal Care and Use Committee of the Institute of Neuroscience, Chinese Academy of Sciences (Center for Excellence in Brain Science and Intelligence Technology, Chinese Academy of Sciences). Male mice, 2- to 3-month-old, D1-Cre (MMRRC_030989-UCD) and D2-Cre (MMRRC_032108-UCD) backcrossed into Black C57BL/6J were used. Male mice, 2- to 3-month-old Black C57BL/6J and PV-Cre mice (Jax stock #008069) were also used. Mice had no previous history of any other experiments. Mice used for behavioral experiments were housed in a room with a reverse light/dark cycle. On days without behavioral training or testing, the water-restricted mice received 1 mL of water. On experimental days, mice receive 0.5 to 1 mL of water during each behavioral session which lasted for 1 to 2 hours. Body weight was measured daily and was maintained above 80% of the weight before water restriction.

### Behavioral training

After one-week recovery following surgery, mice were started with water restriction (1 mL per day), which lasted ~7 days before the beginning of behavior training. Mice performed one behavioral training session each day and were allowed to perform the task until sated. Body weights were measured and water consumption was estimated using the body weight difference before and after each session. If mice consumed less than 0.5 mL water per day, additional water supplement was provided.

The behavioral apparatus has been described previously (52, 53). Briefly, experiments were conducted inside custom-designed double-walled sound-attenuating boxes. The mouse auditory decision behavior was controlled by a custom-developed PX-Behavior System. Cosine ramps (5 ms) were applied to the rising and falling of all tone stimuli. The sound waveforms are amplified using ZB1PS (Tucker-Davis Technologies) and delivered by an electrostatic speaker (ES1, Tucker-Davis Technologies) placed on the right side of the animal. The sound system was calibrated using a free-field microphone (Type 4939, Brüel and Kjær) over 3–60 kHz to ensure a flat spectrum (±5 dB SPL).

The behavioral task is based on the auditory-guided two-alternative-forced-choice (2AFC) task on head-fixed mice as described before (52, 53). After mice learnt how to lick left and right water ports, they were trained to discriminate a range of tone frequencies (between 7 and 28 kHz, or between 5 and 20 kHz, distributed in logarithmic scale of octaves) as higher or lower than the defined category boundary, which is the mid line of the logarithmically spaced frequency range, i.e., 14 kHz for the 7-28 kHz range, and 10 kHz for the 5-20 kHz. The frequency ranges were chosen according to the typical hearing range of laboratory mouse (54). Sound intensity was 70 dB SPL for all used frequencies. The initiation of each trial is not explicitly cued, and the animal needs to wait for the tone stimulus to occur. Following the intertrial interval (ITI), a 0.5-1 s random delay was imposed before tone stimulus (duration, 300 ms) to ensure that animals cannot predict the onset of tone stimuli. Mice were required to respond by licking left or right water port within a 3 s answer period following the stimulus offset to report their perceptual judgements of the tone as the high or low frequency categories. Correct responses led to opening of the water valve for a short period of time to dispense a small amount of water reward (~1.5 μL). Error responses lead to a 2~6 s time-out punishment, during which licking to the wrong side would reinitiate the time-out period. To control for systematic side bias, different groups of mice were trained with opposite stimulus-response contingencies, i.e., high-left / low-right, or high-right / low-left (**Figure 1 E,F** and **Supplementary Figure 4**). Mice completed several hundred trials in each behavioral session allowing for sufficient statistical power quantifying perceptual decisions on stimulus categorization. Mice were first trained with the two easy frequencies separated by 2 octaves, e.g., 7 and 28 kHz, or 5 and 20 kHz until reach 85% correct rate (after ~10 session, **Supplementary Figure 1**), after which 6 tone stimuli of intermediate frequencies (between 7 and 28 kHz or 5 and 20 kHz, linearly spaced on the logarithmic scale) were delivered in randomly interleaved trials. For each session, the fraction of trials with tones of the intermediate frequencies is ~30% to keep the performance stable. Psychometric function was obtained by fitting the behavioral data using a 4-parameter sigmoidal function (55, 56)

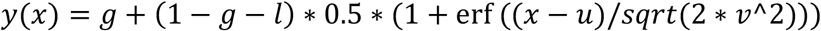

where *y*(*x*) is the probability that animal would make a right choice (**Figure 1C**) or a contralateral choice, and *x* is the tone frequency (in octave). Parameters to be fitted are: *g* the guess rate, *I* the lapse rate, *u* the subject bias (boundary), and *v* the discrimination sensitivity (threshold). erf () represents error function. For psychometric function fitting, ‘miss’ trials in which mice did not response during the answer period were excluded. To quantify the proportion of contralateral choices in optogenetic manipulation and imaging experiments, we pooled all animals trained with either of the two types of contingencies (‘low-left / high-right’ or ‘low-right / high-left’) by ordering the tone frequencies according to relative octaves distance from the boundary frequency, with negative values representing frequencies associated with ipsilateral choices and positive values the contralateral choices.

### Surgery

Mice were anesthetized with isoflurane (1~2%) mixed in air (Pin-Indexed fill or Funnel-Fill Isoflurane Vaporizer, VETEQUIP), and placed on a stereotaxic frame (RWD 68043 and 68077). Throughout the surgery, body temperature was monitored and maintained at 37 °C by temperature controller (RWD 69002), and eyes were moisturized with eye ointment. After shaving the hair, the skin was sterilized with 75% ethanol and iodine. A piece of skin was removed to expose the skull. After cleaning the skull with a scalpel removing connective tissues, a small craniotomy (around 0.5 mm in diameter) was made on the skull above the target area for virus injection. Virus was injected slowly with a hydraulic manipulator (Narashige, MO-10) through a glass micropipette (pulled using Sutter Instrument P-97, with tip broken into an opening with 20-30 μm inner diameter; Drummond Scientific, Wiretrol II Capillary Microdispenser) into the target area. After injection, the micropipette was kept in the injection site for ~10 min before being retracted slowly to reduce flow back of the viruses. Virus preparations were obtained from Shanghai Taitool Bioscience Co.Ltd, or OBiO Technology (Shanghai) Corp., Ltd. About 30 min after virus injection, fiber optics for fiber photometry recording or optogenetic manipulation (NA 0.37, 200 μm core, ceramic ferrule OD 1.25mm) were implanted ~200 μm above the virus injection sites. After injection of virus and implantation the fiber optics, silicon adhesive (Kwik-sil, WPI) was used to cover the craniotomy. The exposed skull and surrounding tissues were then covered with instant adhesive (LOCTITE 495, Henkel). A custom-designed titanium headplate was attached to the skull with dental cement for head fixation. After surgery, mice were allowed to recover at least 7 days before water restriction and behavioral training.

For non-selective inactivation of striatal neurons, 400 nL AAV2/9-CAG-ArchT-GFP (3.50×1012 V.G./mL) virus was injected to TS (AP: −1.46, ML: 3.25, DV: −3.4) or to the anterior sub-region of dorsal stratum (AP: −0.02, ML: 2.0, DV: −3.4) in wildtype C57BL/6J mice (**Figure 1**). For control animals, AAV2/9-CAG-eGFP-2A (2.40×1013 V.G./mL) virus was injected into TS (**Figure 1 H,I**).

For optogenetic manipulations of specific types of striatal neurons, 400 nL AAV2/8-CAG-FLEX-ChR2(H134R)-mCherry (for activation, Figure 2-figure supplement 1, 2.58×10^13^ V.G./mL) or 400 nL AAV2/9-Ef1α-DIO-eNpHR3.0-eYFP (for inactivation, **Supplementary Figure 7** and **Supplementary Figure 9A**, 1.50×10^13^ V.G./mL) virus was slowly injected into TS in D1-Cre, D2-Cre or PV-Cre mice (**Figure 2; Figure 3; Figure 4 A,B**). For iSPN activation by ChrimsonR or iSPN inactivation by stGtACR2, AAV2/9-hSyn-FLEX-ChrimsonR-tdTomato (**Figure 5A-C**) or AAV2/9-hSyn-SIO-stGtACR2-FusionRed (**Figure 5D-F**) was injected into TS in D2-Cre mice.

For fiber photometry recording from the PV neurons, 300 nL AAV2/9-hSyn-FLEX-GCaMP6s was injected into bilateral TS of PV-Cre mice, and fiber optics (NA 0.37, 200 μm core, ferrule O.D. 1.25mm) were inserted ~200 μm above the virus injection sites (**Figure 4D-F**). For fiber photometry recording from dSPNs axons in SNr, 300 nL AAV2/9-hSyn-GCaMP6s was slowly injected in bilateral TS (AP: −1.46, ML: 3.25, DV: −3.4) of D2-Cre mice, and fiber optics (NA 0.37, 200 μm core, ferrule O.D., 1.25mm) were inserted into the bilateral SNr (AP: − 3.5, ML: 1.8, DV: 4.3) (**Figure 5**).

### Optogenetic manipulations

For cell-type specific activation by ChR2, a 473-nm diode-pumped solid state laser (BL473T8-100FC, Shanghai Laser & Optics Century Co., Ltd.) with the output power adjusted to ~2 mW at the distal end of patch cable was used to activate ChR2-expressing neurons. For cell-type specific inactivation by NpHR3.0, a 593-nm diode-pumped solid state laser (YL593T6-100FC, Shanghai Laser & Optics Century Co., Ltd.) with output power of ~20 mW was used to inactivate NpHR-expressing neurons. For iSPN inactivation by stGtACR2, a 462-nm blue diode laser (BLM462TA-300FC, Shanghai Laser & Optics Century Co., Ltd.) with an output power of 2 mW was used to inactivate stGtACR2-expressing iSPNs. For iSPN activation by ChrimsonR, a 635-nm red diode laser (RLM635TA-100FC, Shanghai Laser & Optics Century Co., Ltd.) with an output power of 5 mW at the distal end of patch cable was used to activate ChrimsonR-expressing iSPNs. For inactivation experiments using ArchT and control group expressing GFP, a 532-nm diode-pumped solid state laser (GL532TA-100FC, Shanghai Laser & Optics Century Co., Ltd.) with an output power of 20 mW at the distal end of patch cable was used.

In all optogenetic stimulation sessions, photostimulation was applied to a proportion of trials randomly interleaved with control trials (no photostimulation) to allow for within-subject and within-session comparison. Since in most sessions there were less trials with intermediate frequencies (~30%) and more trials with the end frequencies (~70%), to obtain comparable number of trials with photostimulation for intermediate and end frequencies, photostimulation was applied to ~50% of the intermediate-frequency trials, and ~15% of end-frequency trials.

### Fiber photometry recording

The fiber photometry system was custom built based on previous method (57). The excitation light was from a 470-nm LED (Cree, Inc, XQEBLU-SB-0000-00000000Y01-ND) and a 410-nm LED (Xuming, 3W, 410-420 nm). The intensity of the 470-nm light was set to 10-15 μW measured at the distal end of the patch cord (calibrated based on the transmittance of the fiber optics implanted in the brain, to keep the intensity at 10 μW around the fiber optic tips). For each recording site, the intensity of 410-nm light was adjusted to keep the recorded emission fluorescence level excited by 410-nm light close to that excited by 470-nm light from the brain tissue.

The lights from 470-nm LED and 410-nm LED independently passing through aspheric condenser lenses (Thorlabs, ACL25416), were filtered through respective bandpass filters (FB470-10, and FB410-10, Thorlabs), and were then combined by a dichroic mirror (DMLP425R, Thorlabs). The combined excitation lights then sequentially passed through a dichroic mirror (Chroma, T525/50dcrb) and an objective (Olympus, UPLFLN 20X NA0.5), and then were coupled into a fiber optic cable (NA 0.37, 200 μm core) that affixed to the fiber optics implanted in mouse brain. A small fraction of 470-nm and 410-nm lights reflected by the first dichroic was collected by a condenser lens (Thorlabs, ACL25416) and received by a photodiode (Thorlabs, PAD100A2), the real-time light intensity of which was used as a feedback signal for custom-designed closed-loop circuits to stabilize the intensity of 470-nm and 410-nm lights throughout the recording session (**Figure 4D; Supplementary Figure 11**).

The emitted fluorescent signals were collected by the same light path and focused onto a CCD (QImaging, R1 retina). The 470-nm LED, the 410-nm LED and the CCD were controlled by a NI I/O device (NI, USB-6341) and software written in LabVIEW. The NI device also receives the triggers from the digital outputs of PX-Behavior System to tag the behavior-related time. Consecutive camera frames were captured at 40 Hz using alternating 470-nm and 410-nm excitation lights (20 Hz sampling rate for both 470 nm and 410 nm channels). The excitation pulse duration of 470-nm LED or 410-nm LED in each frame was 11 ms for an exposure time of 10 ms, beginning at 0.5 ms prior to the onset of the exposure time of CCD and ending at 0.5 ms after the offset of the exposure time of CCD (**Supplementary Figure 11B**). The signals from CCD were recorded by ImageJ based software Micro-Manager (Version 2.0 beta), with the ‘binning’ at 16×16, and the ‘Exposure’ time of 10ms. The fluorescent signals excited by the 470-nm and 410-nm were extracted and analyzed subsequently using MATLAB scripts.

For simultaneous activation of iSPNs (expressing ChrimsonR) and fiber photometry recording from dSPN axons (**Figure 5A-C**), 1 s of 635-nm continuous red light was delivered from sound onset in photostimulation trials, and the settings of fiber photometry recording were same as those used in **Fig. 4** (**Supplementary Figure 11B**). For simultaneous inactivation of iSPNs (expressing stGtACR2) and fiber photometry recording from dSPN axons (**Figure 5D-F**), the laser pulses (462 nm) for optogenetic stimulation and the fluorescence signal acquisition were alternating in time to avoid potential contamination from 462 nm laser to fluorescence signals. At each sampling point, the fluorescence signals were excited by 5 ms 470 nm (~30 uW) or 410 nm, alternating frame by frame, and collected by CCD with 4 ms exposure time within the excitation period. The fluorescence excitation was then followed by a 20 ms optogenetic laser pulse (~ 4 mW, 462 nm) (**Supplementary Figure 11C**). Each photostimulation/recording cycle was repeated at 40 Hz.

### Optogenetics data analysis

For analyzing the behavioral effects of optogenetic stimulation, ‘miss’ trials were excluded. To quantify the behavioral effects of optogenetic stimulation, we used the two-way repeated-measures ANOVA with function ‘fitrm’ in Matlab to test the effect of optogenetic manipulation across tone frequencies and used LSD post hoc tests to compare the effects at each frequency.

### Fiber photometry data analysis

The 410-nm reference trace was scaled to best fit the 470-nm signals using least-squares regression (58). Then the motion-corrected 470-nm signal (F_470_) was obtained by subtracting the scaled 410-nm reference trace from the 470-nm signal and then adding the mean value of scaled 410-nm trace. ΔF/F signal was calculated by subtracting the mode of F_470_ and then dividing by the mode value of F_470_. To compare signals across recording sessions, we used the z-score of calcium signals, which is the ΔF/F subtracted by the mean and divided by the standard deviation of the data in the session. The peak value of z-score within 1 s after answer time was used for statistics in **Figure 4F**, and the peak value of z-score within 1 s after sound onset (optogenetic period) was used for comparison in **Figure 5C,F**.

### Statistical Analysis

All statistical analysis was performed using MATLAB 2017a (MathWorks). All data are shown as mean ± s.e.m. unless mentioned otherwise. The comparison significance was tested using paired two-sided Wilcoxon signed rank test, two-way repeated-measures ANOVA and post hoc Fisher’s LSD multiple comparison tests. *P < 0.05, **P < 0.01, ***P < 0.001. For activation of iSPNs, one session was excluded due to inconsistent sound frequency setting with other sessions.

### Histology

After the completion of experiments, mice were deeply anesthetized and perfused with saline and 4% PFA (or 10% formalin), and brains were removed and post-fixed overnight. Brains were then incubated with PBS containing 30% sucrose overnight. Frozen sections (50 μm, using Leica CM1950 cryostats) containing viral injection sites, cannula window location and optic fiber tracks were collected and stained with DAPI to visualize cell nuclei. Histology images were taken using Olympus VS120® Virtual Microscopy. For verification of cell-type specific virus expression in dSPNs and iSPNs, 200 nL AAV2/8-CAG-FLEX-eGFP-WPRE-pA (1.25×10^13^ V.G./mL, TaiTool Bioscience) was injected into the TS of D1-Cre or D2-Cre mice. One month after the surgery, the animals were perfused and the brains were sectioned sagittally to examine the projection pattern of neurons expressing GFP (**Supplementary Figure 3A-B**).

### Neural network model

To examine the potential computational implementation for how concurrent but asymmetrical dSPN and iSPN activity could result in unambiguous action selection following decisions, we constructed a computational model of the cortico-basal ganglia circuitry on the population level, which is similar to the computational model of dopamine-biasing action selection through basal ganglia in the previous study (59). In the model, the decision outcomes are transferred from cortex to dSPNs, iSPNs and striatal PV neurons via canonical cortico-striatal projections (60). The activity of the dSPNs and iSPNs populations converge to the ipsilateral substantia nigra pars reticulata (SNr) populations through the direct and the indirect pathway respectively. The left and right SNr populations mutually inhibit each other, and the dominant SNr population determines the motor output by controlling the final motor output either through brainstem circuits or motor cortices. The network model composed of neural populations in both left and right hemispheres with symmetrical connectivity, as shown in **Figure 6**.

In the model, a perceptual decision *d* is produced in the cortex according to a simple decision rule (Equation 1). Where *sign*(*q*) = 1 if *q* ≥ 0, *sign*(*q*) = 1 if *q* < 0. Note that the tone of stimulus *q* ∈ [−1,1] is in the unit of octaves. *θ*(*q*) is the probability of wrong decision due to perceptual uncertainty, where *σ* = 2.24 is the degree of uncertainty in the perception (Equation 2). The decision outcome *d* = ±1 encodes left or right choice. The cortical neurons in each hemispace are organized into two types of populations that are selective to contralateral or ipsilateral choices. The firing rate of cortical population *CTX_jk_* is shown in Equation 3, where *j* ∈ {*L, R*} representing the left or right hemisphere, *k* ∈ {*contra, ipsi*} is the selectivity for left (*s* = −1) or right (*s* = 1) action, *s* = –1 corresponds to *f^CTX_Rcontra_^* and *f^CTX_Lipsi_^*, *s* = 1 corresponds to *f^CTX_Lcontra_^*, and *f^CTX_Ripsi_^*, and *d* is the decision outcome given by Equation 1. *I* = 20 is the peak firing rate of cortical neurons. *f_b_* = 0 is reference frequency. As the frequency *q* changes, the firing activity of cortical neural population switches between low and high activity states in a sigmoid form, with the switching slope controlled by ζ = 0.01. Cortical neurons keep self-sustained activity states for a period of time without external input (61).

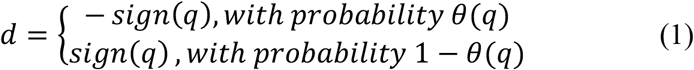

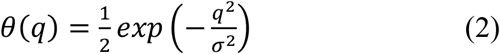

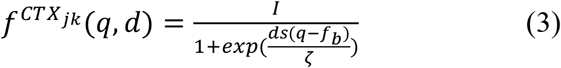

SPNs in the striatum receive currents both externally and internally (Equation 4), where 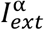 and 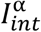 are the external current and internal current to population *α*, from the cortex and from striatal sub-populations respectively (**Figure 6A**). SPNs are divided into 8 sub-populations based on different response preferences found in our imaging results and from the two hemispheres, *α* = *D_ijk_*, with *i* ∈ {1,2} representing cell types (dSPNs and iSPNs). The cortical input to population α is shown in Equation 5, where *f^CTX_jk_^* is the activity of the cortical population with selectivity *s* (Equation 3). 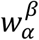 is the connection strength from population β to population α (**Supplementary Table 1**). *V_α_* is the membrane potential. For the excitatory connection from *CTX_jk_*, the reverse potential *E_CTX_jk→α__* = 0. The internal input current to population α is shown in Equation 6, where *β* is a striatal population including α and PV, and *f*^β^ is its firing activity. Reverse potential parameters are set according to precious study (62). For inhibitory connections from striatal population *β*, the reverse potential was *E_β→α_* = −65*mV*. For the connection from iSPN to SNr, *E*_*D*_2*jk*_→*SNr_j_*_ = 0 *mV*.

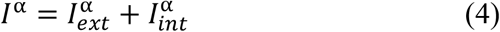

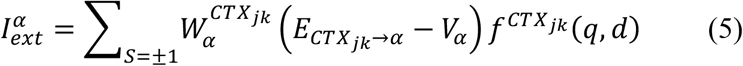

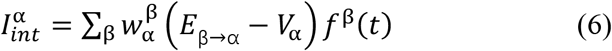

Take *D*_1,*L,Contra*_ as an example of SPN sub-population to explain its specific input current shown in Equation 4–6,

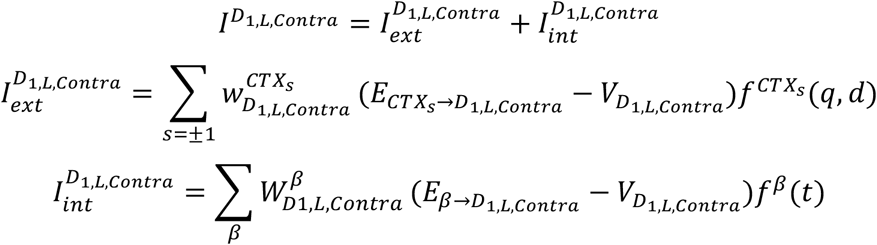

PV neurons only receive cortical inputs from ipsilateral cortex, 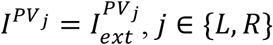, take PV neurons in the left hemisphere as an example,

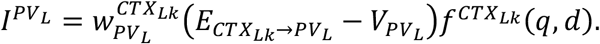

SNr neurons receive external current from SPNs and internal current as their inputs. Take SNr neurons in the left hemisphere as an example,

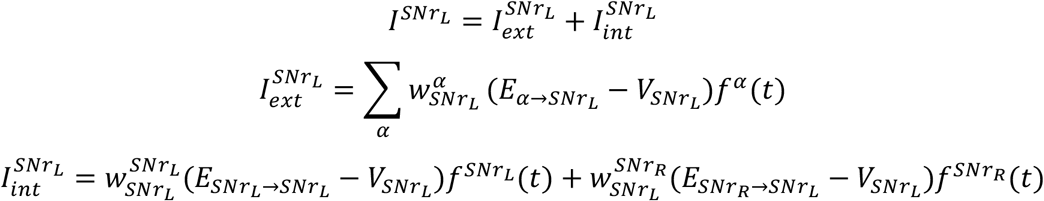

The membrane potential of a striatal population α is described as shown in Equation 7. *E_a_* = −55 *mV* is the reversal potential. ϵ is a noise term. The firing frequency of a striatal population α is shown in Equation 8. The transfer function *g* is linear above the discharge threshold θ = −40 *mV* and has an exponential growth trend below the discharge threshold (Equation 9), where *k* = 1 represents medium spiny projection neurons, and *k* = 5 represents interneurons in the striatum.

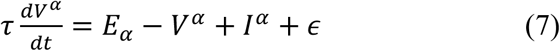

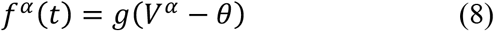

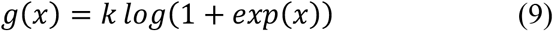

The motor choice is made according to the SNr activity after 100 ms when the network activity becomes stabilized. The rule for motor choice is defined in Equation 10.

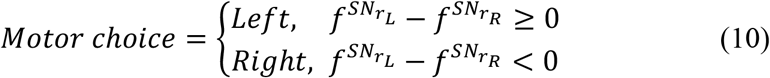

The connection weights between the populations in the network are shown in **Supplementary Table 1**. The weights are assumed to form through pervious learning processes. Populations are labeled according to the convention introduced in previous sections. For example, *CTX*_−1_ is the cortical population selective to left choice, and *D*_1*Rcontra*_ is the dSPN population connected to the right SNr and is selective for contralateral choice. For simplicity, we omitted some indices of the labels if they are not concerned. For example, *D*_1*L*_ includes populations *D*_1*Lipsi*_ and *D*_1*Lcontra*_. The dSPNs and iSPNs in the striatum inhibit each other, and the weights are shown as 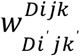.

The optogenetic manipulation of a striatal population *α* is modelled by an additional current *I^stim^* to the membrane potential dynamics (Equation 11), where *I^stim^* is the regulation current, taking positive or negative values for optogenetic activation or inactivation, respectively. The parameters of optogenetic regulation are shown in **Supplementary Table 2**.

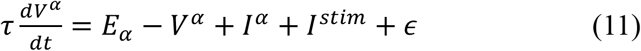

We performed the network model stimulation using the same set of parameters under the standard control conditions (**Supplementary Table 1**), as well as under the perturbed conditions caused by optogenetic activation or inactivation (**Supplementary Table 2**). Under each condition, 300 independent trials were simulated, and the probability and the standard deviation of choosing right choice was calculated. Psychometric functions were fitted by a 4-parameter sigmoidal curve same as in the experimental data. Compared with the standard control situation, under the perturbed condition of optogenetic activation/inactivation of striatal PV neurons, dSPNs and iSPNs, the model reproduced similar biased action selection behavior as observed under the corresponding conditions in experiments (**Figure 6C-G**). The psychometric functions of action selection of the model qualitatively captured the deviation from that of the control condition after perturbation as shown in experiments.

## Acknowledgements

We thank Dr. Mu-ming Poo, Dr. Yong Gu, Dr. Yangang Sun and Dr. Chengyu Li for valuable discussions; Dr. Chunyu Ann Duan and Dr. Yu Xin for help in data analysis; Mengjun Sheng and Di Lu for help in surgery; Zhaomei Ying for lab assistant.

## Author contributions

L.C., S.T. and N.L.X. conceived the project and designed the experiments. L.C. performed the optogenetic experiments. S.T. developed striatum imaging method and performed fiber photometry recordings. J.P. designed the fiber photometry system. J.P. and N.L.X. developed the behavioral system. K.Z. and B.S. constructed the neural network model and performed the model simulation. Z.Z. performed part of the behavioral training and optogenetic experiments. L.C., S.T. and N.L.X. analyzed the data and wrote the manuscript.

## Materials & Correspondence

Correspondence and requests for materials should be addressed to N.L.X.

## Data and materials availability

All data to understand and assess the conclusions of this study are available in the main text or supplementary materials. All the original behavioral, imaging, histochemical data and analysis code are archived in the Institute of Neuroscience, CAS Center for Excellence in Brain Science and Intelligence Technology, Chinese Academy of Sciences.

---

**Supplementary Figure 1.**
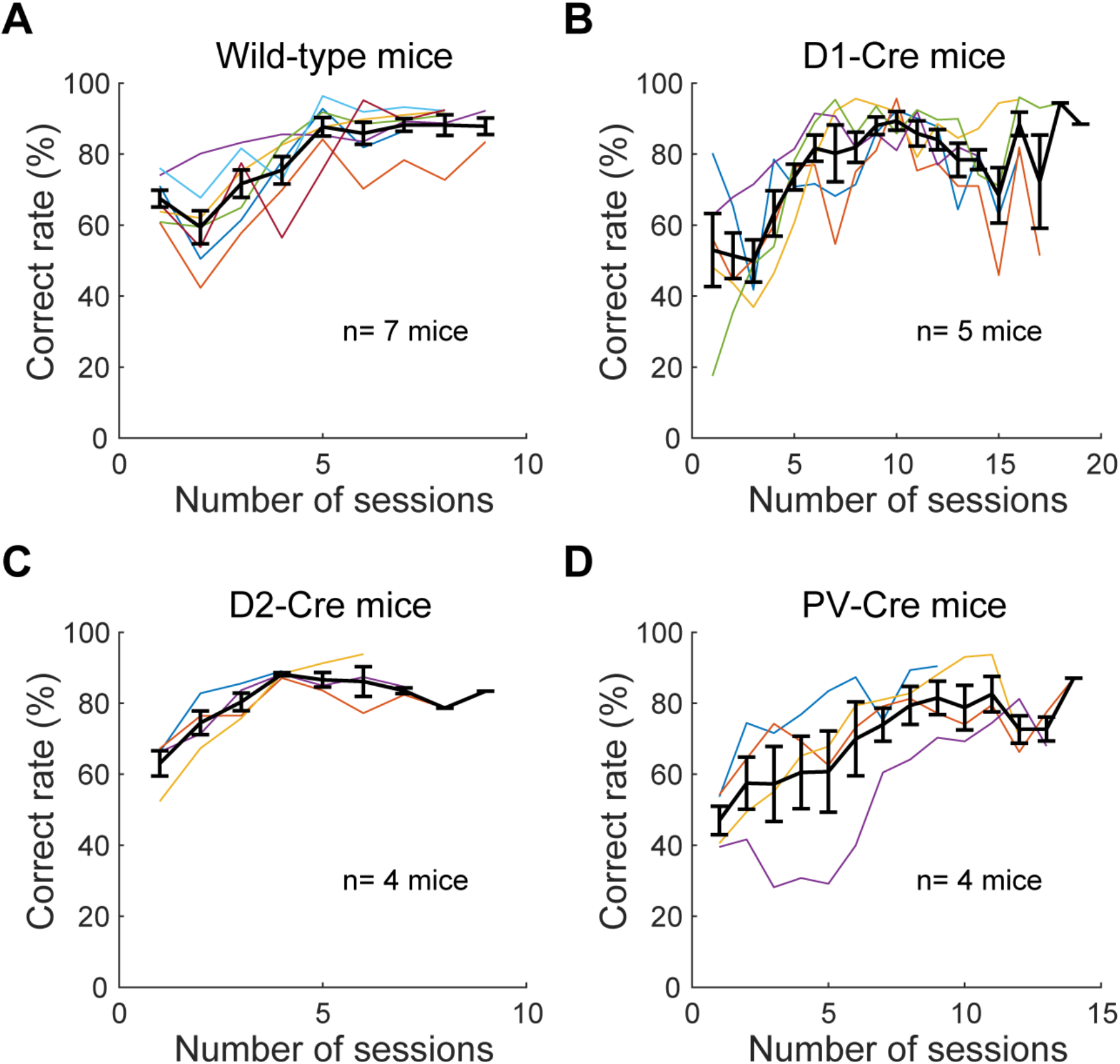
Learning curves of example mice. (A) The learning curves of example wildtype mice (n= 7 mice). (B) The learning curves of example D1-Cre mice (n=5 mice). (C) The learning curves of example D2-Cre mice (n= 4 mice). (D) The learning curves of example PV-Cre mice (n= 4 mice). Thin lines are correct reate form individual mice. Black lines and dots are shown as mean ± s.e.m. for each group of animals.

**Supplementary Figure 2.**
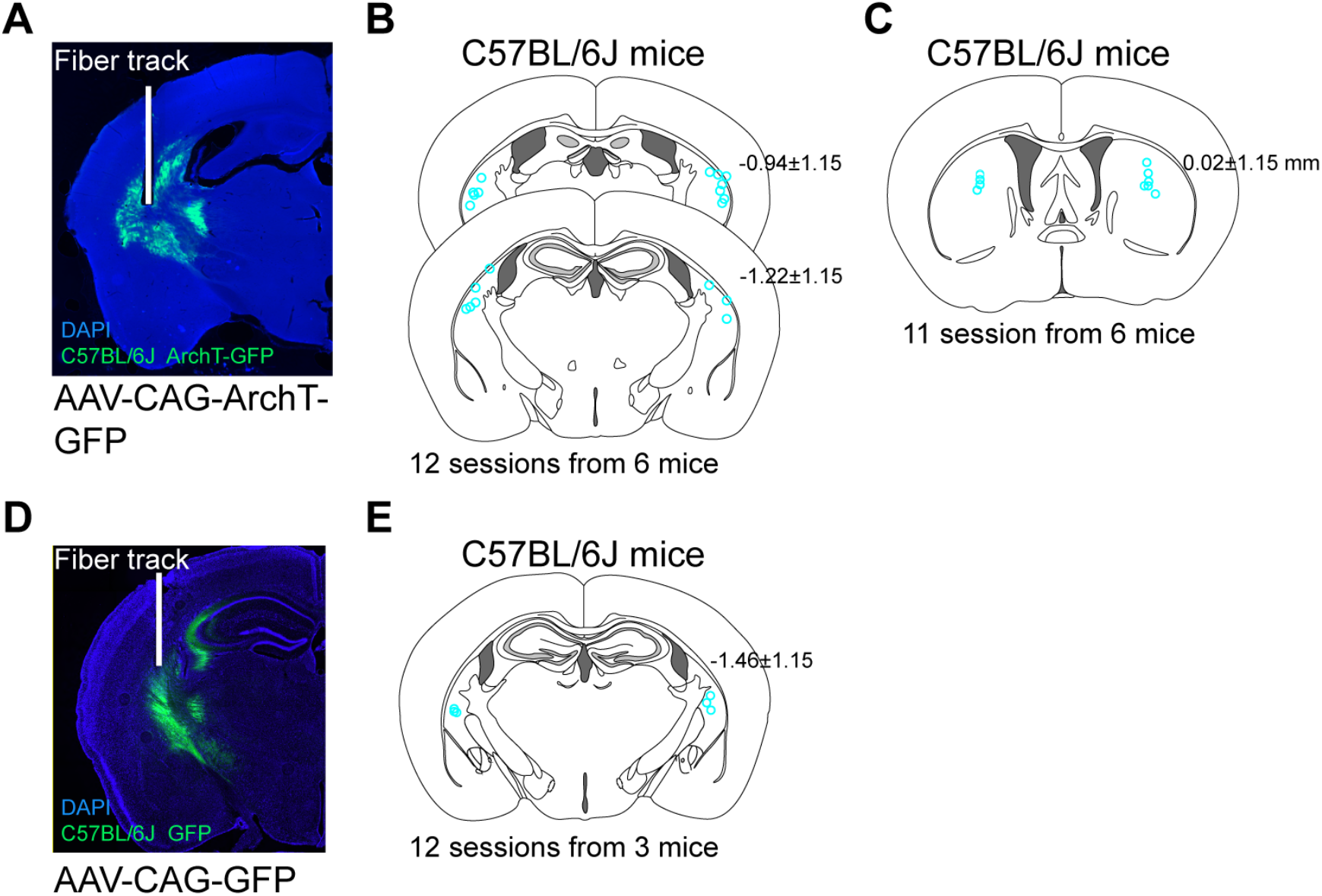
Histological images of virus injection and fiber track for optogenetic inactivation. (A) Example histological image of a C57BL/6J mouse injected with AAV-CAG-ArchT-GFP in the TS. (B) The locations of fiber tips (cyan circles) for optogenetic inactivation of the TS by ArchT. (C) The locations of fiber tips (cyan circles) for optogenetic inactivation of the anterior sub-region of the dorsal striatum by ArchT. (D) Example histological image of a C57BL/6J mouse injected with AAV-CAG-GFP in the TS. (E) Locations of fiber tips (cyan circles) for mice expressing GFP in the TS.

**Supplementary Figure 3.**
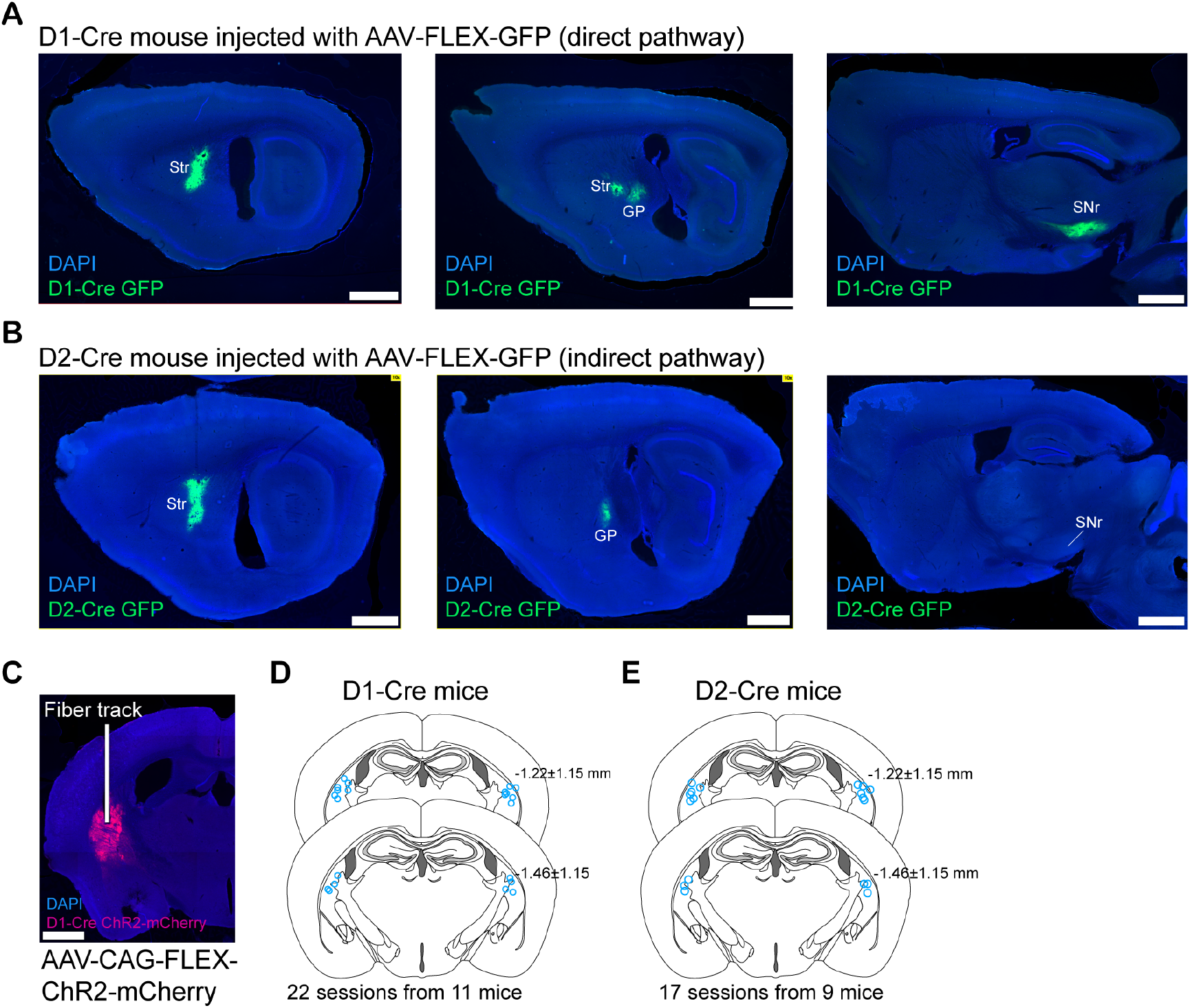
Histology verifying the cell type specific virus labeling of SPNs and the optogenetic activation locations. (A) Sagittal section from an example D1-Cre mouse, showing Cre-dependent viral expression (GFP) in the TS and downstream axonal targets (GP and SNr) of labeled dSPNs. Scale bars, 1 mm. (B) Sagittal section from an example D2-Cre mouse, showing Cre-dependent viral expression (GFP) in the TS and downstream axonal targets (GP only) of labeled neurons. (C) Example histological image of a D1-Cre mouse injected with AAV-FLEX-ChR2-mCherry in the TS. (D) Locations of fiber tips (blue circles) for optogenetic activation in D1-Cre mice by ChR2. (E) Locations of fiber tips (blue circles) for optogenetic activation in D2-Cre mice by ChR2.

**Supplementary Figure 4.**
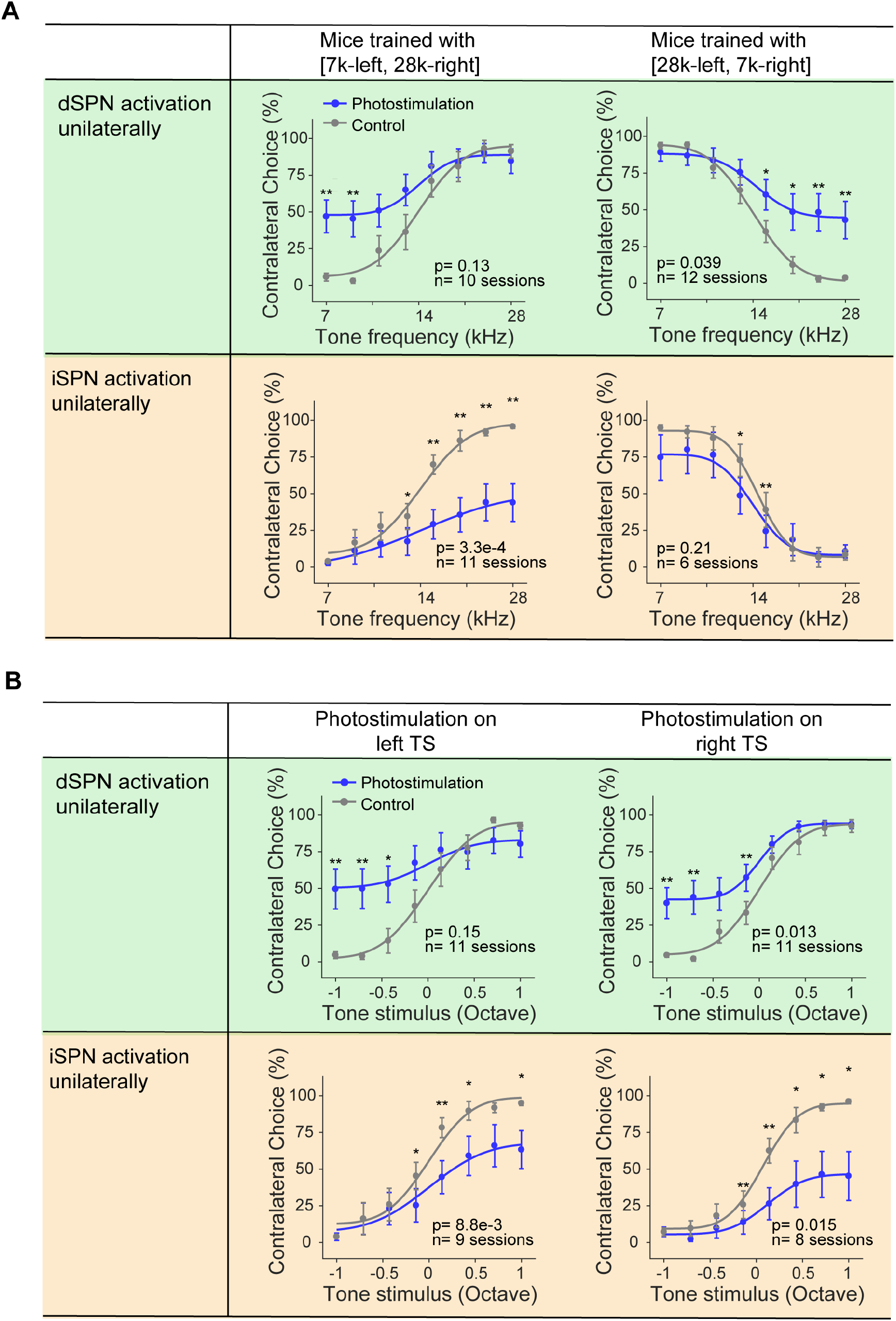
The effects of optogenetic activation with different sound-choice association and in left/right hemisphere respectively. (A) Psychometric functions from the two groups of mice trained with reversed stimulus-choice associated were summarized separately, showing percentages of choices contralateral to the photostimulated hemisphere as a function of the tone frequency values. Top two are from dSPN activation, and bottom two are from iSPN activation. (B) Summarized psychometric functions from control trials (grey) and trials in the left TS and in the right TS, showing percentages of choices contralateral to the photostimulated hemisphere as a function of the tone stimulus. The tone stimuli were arranged such that the positive values indicate tones associated with contralateral choices and negative values associated with ipsilateral choices. Top two are from dSPN activation, and bottom two are from iSPN activation. Two-way repeated-measures ANOVA; *p < 0.05, **p < 0.01, Fisher’s LSD post hoc tests.

**Supplementary Figure 5.**
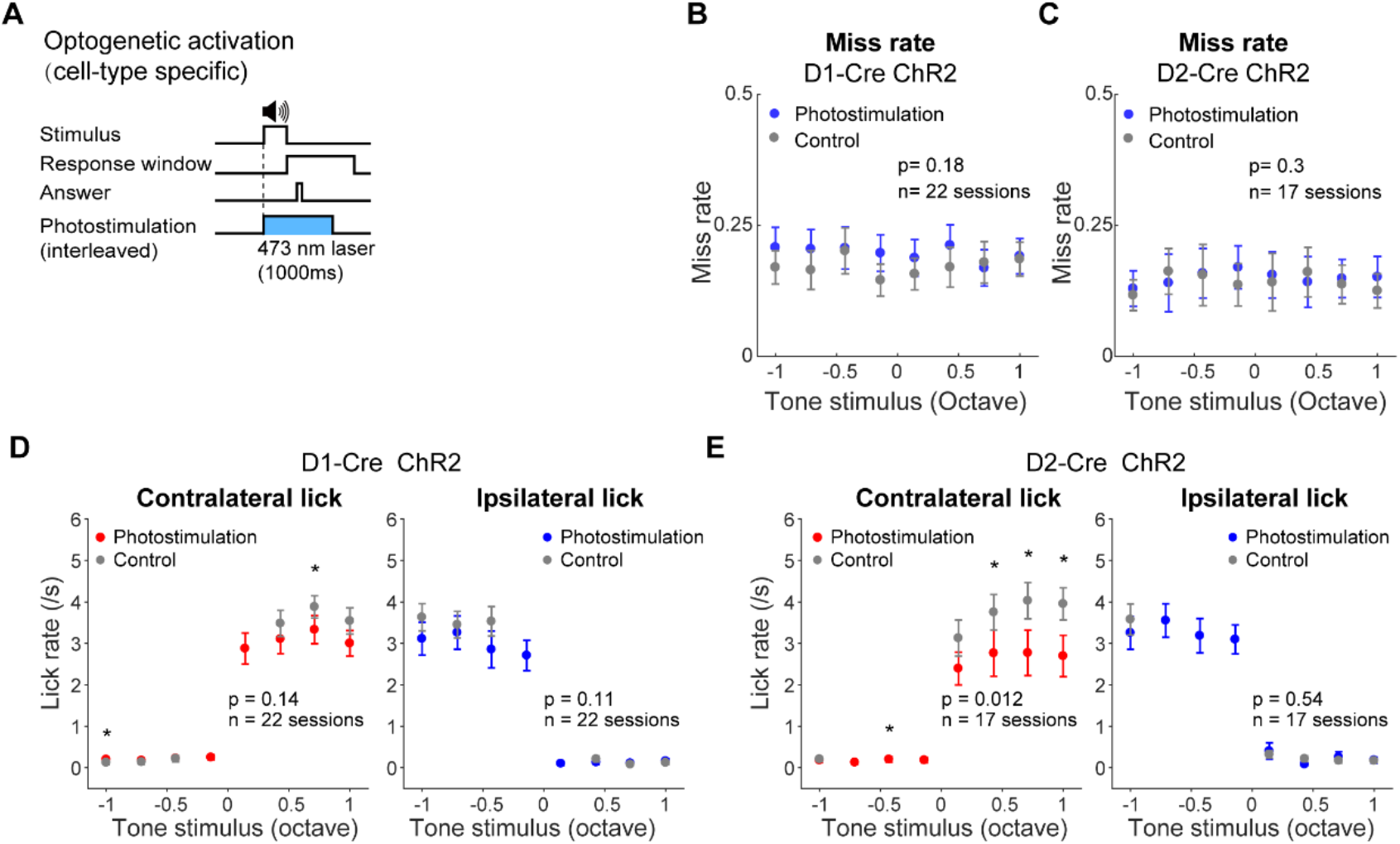
The effects of cell-type specific optogenetic activation on miss rate and lick rate. (A) Schematic showing trial structure with optogenetic activation by ChR2. (B-C) Miss rate as a function of tone stimulus in trials with or without optogenetic activation of dSPNs (B) or activation of iSPNs (C). (D-E) Lick rate as a function of tone stimulus in trials with or without optogenetic activation of dSPNs (D) or activation of iSPNs (E).Left, effect on contralateral licks; right, effect on ipsilateral licks. Two-way repeated-measures ANOVA; *p < 0.05, Fisher’s LSD post hoc tests.

**Supplementary Figure 6.**
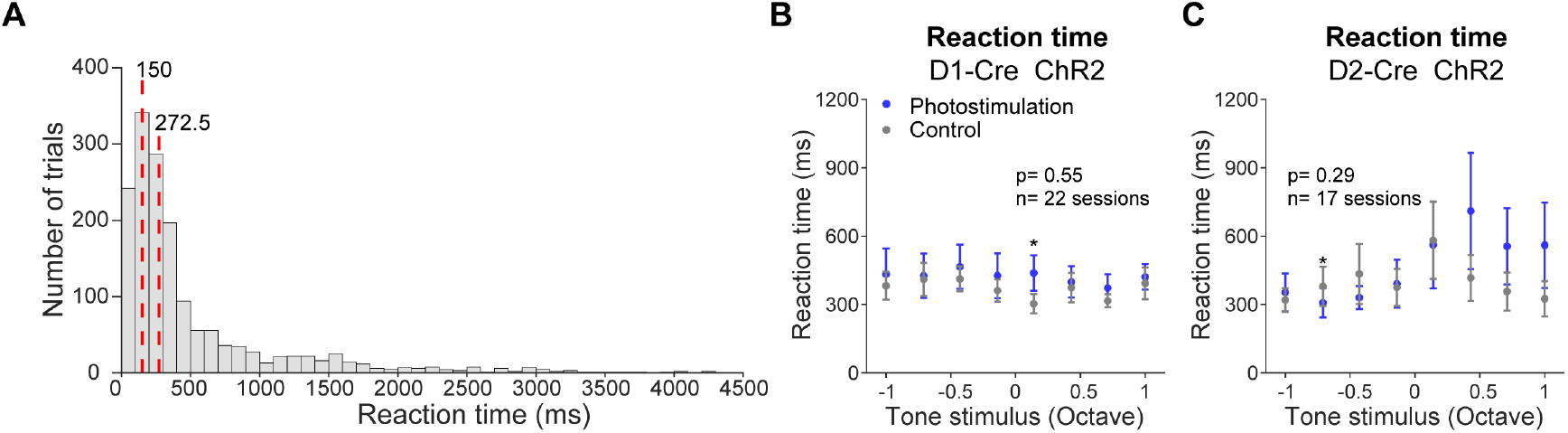
Reaction time of mice in the auditory-guided decision making. **(A)** Distribution of reaction time in control trials for D2-Cre mice as shown in Figure 2E. Vertical red lines indicate the first quarter (150 ms) and the median (272.5 ms). **(B-C)** Reaction time as a function of tone stimulus in trials with or without optogenetic activation of dSPNs (**B**) and activation of iSPNs (**C**), respectively. Twoway repeated-measures ANOVA; *p < 0.05, Fisher’s LSD post hoc tests.

**Supplementary Figure 7.**
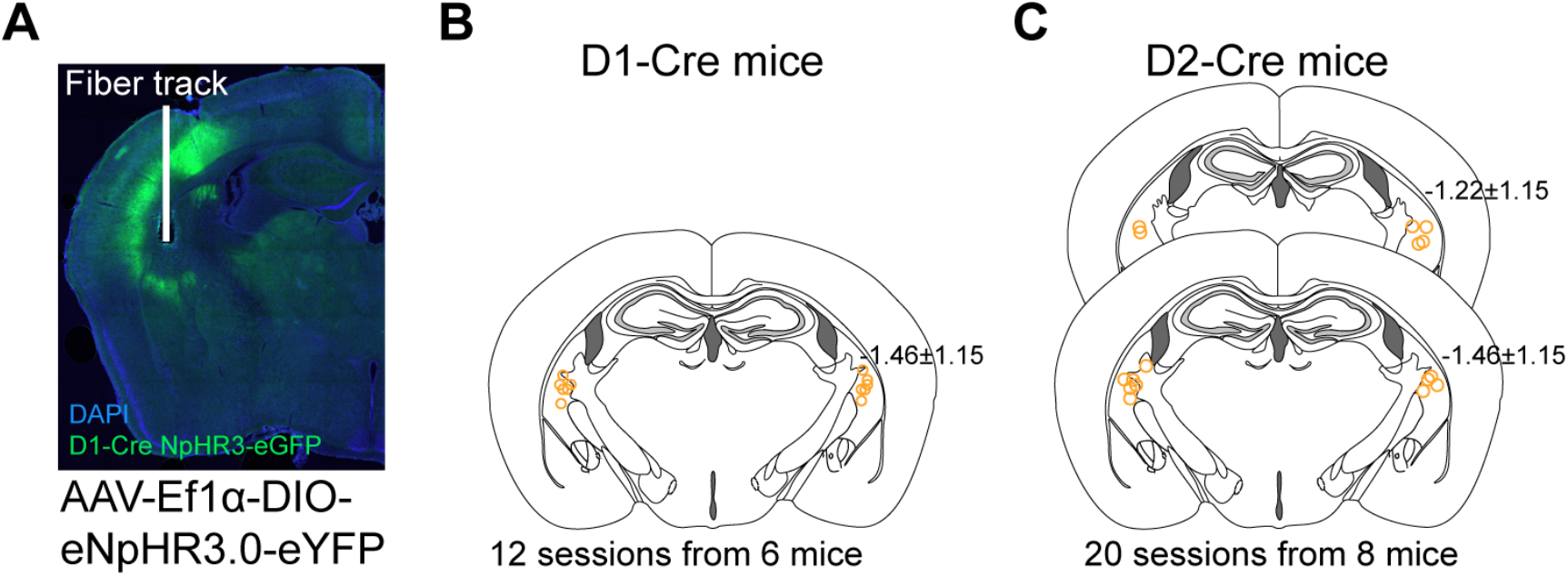
Histological images of virus injection and fiber track for optogenetic inactivation of dSPNs and iSPNs. (A) Example histological image of a D1-Cre mouse injected with AAV-DIO-eNpHR3.0-eYFP in the TS. (B) The locations of fiber tips (orange circles) for optogenetic inactivation by eNpHR3.0 in D1-Cre mice. (C) The locations of fiber tips (orange circles) for optogenetic inactivation by eNpHR3.0 in D2-Cre mice.

**Supplementary Figure 8.**
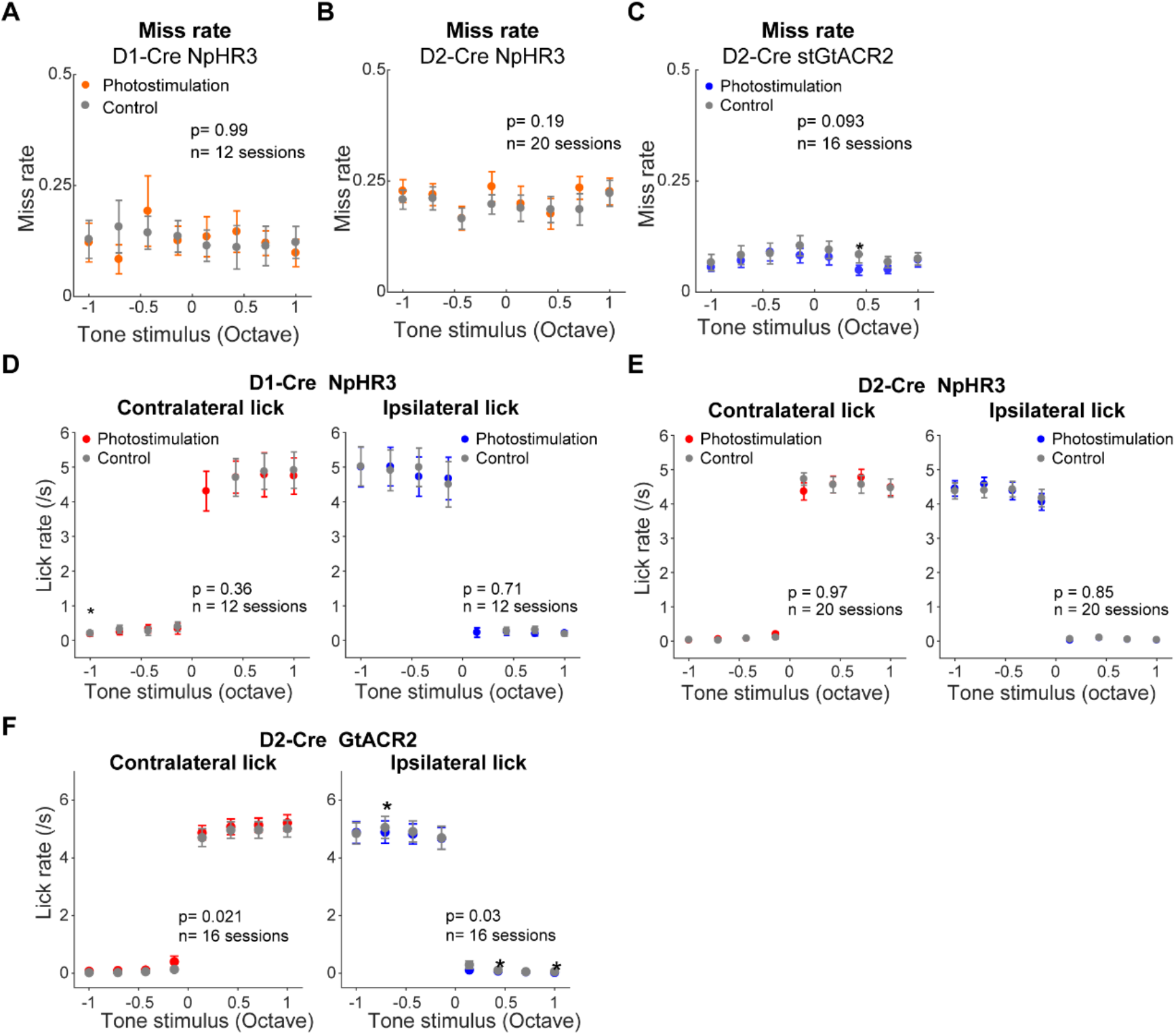
The effects on miss rate and lick rate for optogenetic inactivation of dSPNs and iSPNs. (A-C) Miss rate as a function of tone stimulus in trials with or without optogenetic inactivation of dSPNs by NpHR3 (A), inactivation of iSPNs by NpHR3 (B), and inactivation of iSPNs by stGtACR2 (C). (D-F) Lick rate as a function of tone stimulus in trials with or without optogenetic inactivation of dSPNs by NpHR3 (D), inactivation of iSPNs by NpHR3 (E), and inactivation of iSPNs by stGtACR2 (F). Two-way repeated-measures ANOVA; *p < 0.05, Fisher’s LSD post hoc tests.

**Supplementary Figure 9.**
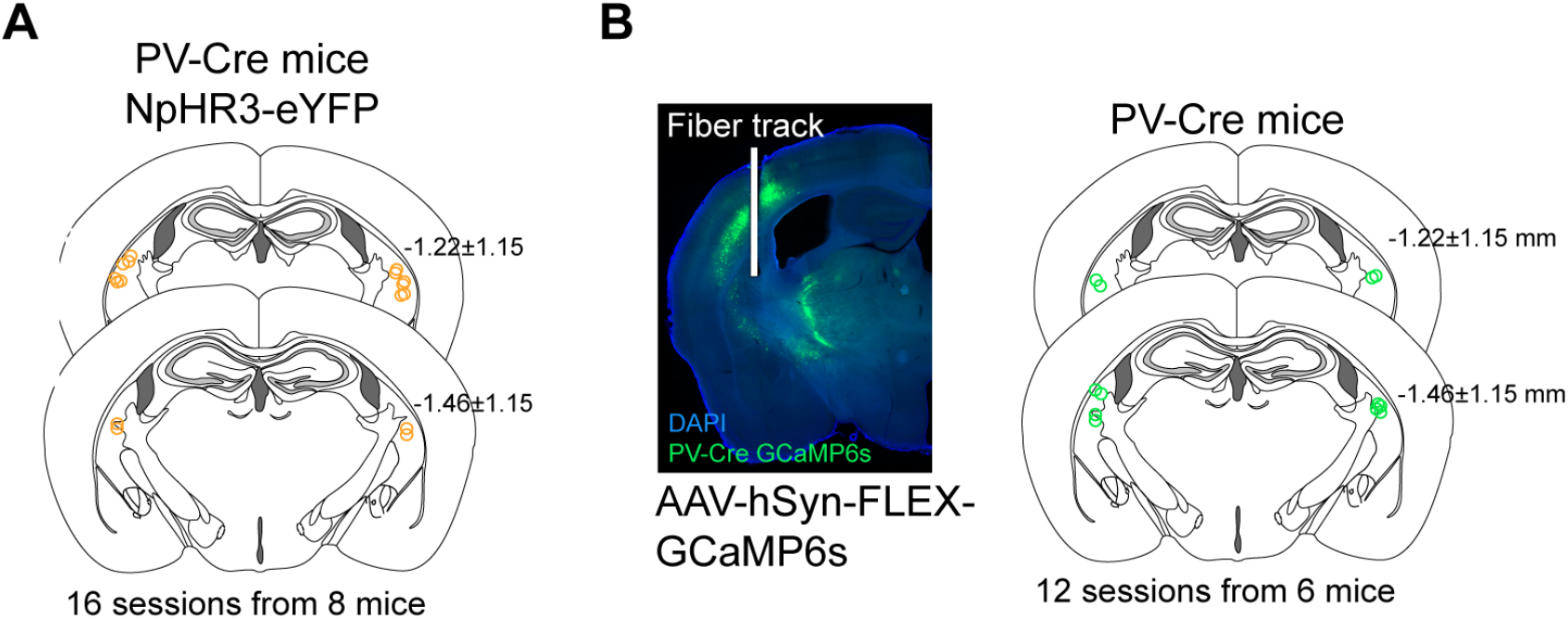
Locations of fiber photometry and optogenetic manipulation for PV interneurons in the TS. (A) The locations of fiber tips (orange circles) for optogenetic inactivation by AAV-DIO-eNpHR3.0-eYFP in the TS of all PV-Cre mice. (B) Left, example histological image of a PV-Cre mouse injected with AAV-FLEX-GCaMP6s in the TS. Right, locations of fiber tips (green circles) in all experiments.

**Supplementary Figure 10.**
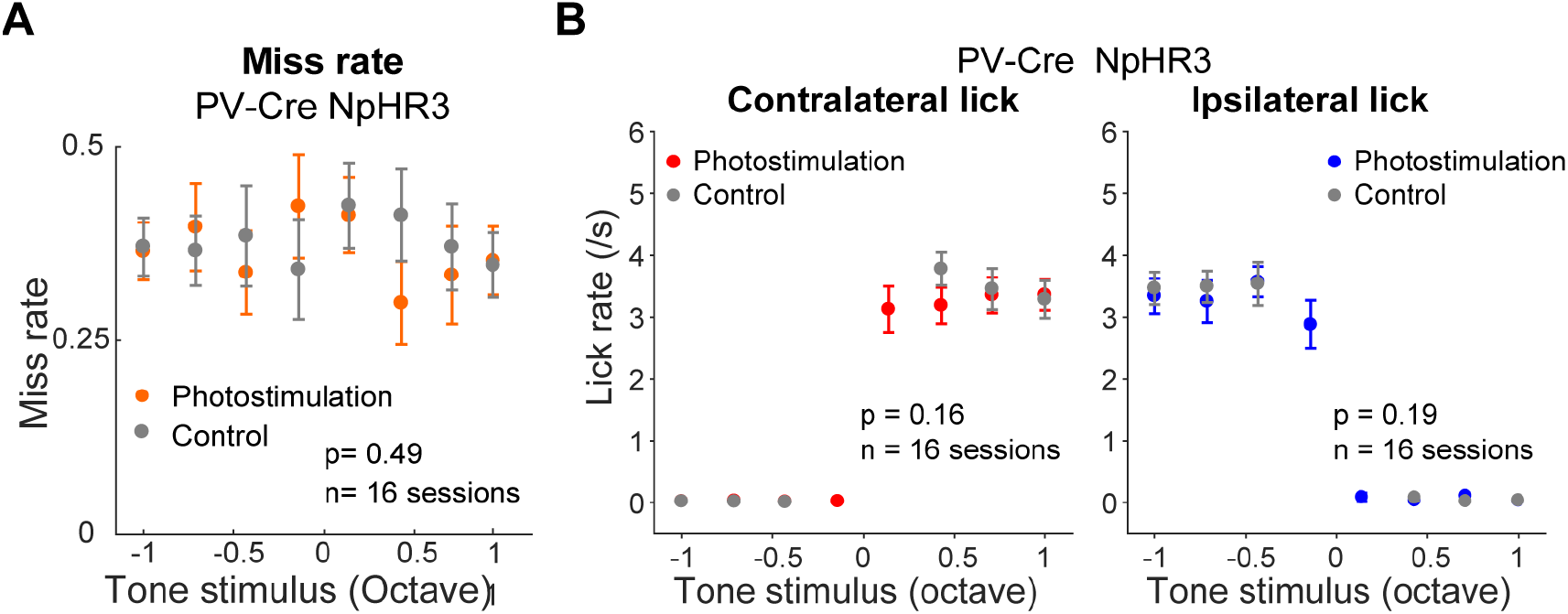
The effects on miss rate and lick rate for optogenetic inactivation of PV interneurons in the TS. (A) Miss rate as a function of tone stimulus in trials with or without optogenetic inactivation of PV interneurons in the TS. (B) Lick rate as a function of tone stimulus in trials with or without optogenetic inactivation of PV interneurons in the TS. Two-way repeated-measures ANOVA.

**Supplementary Figure 11.**
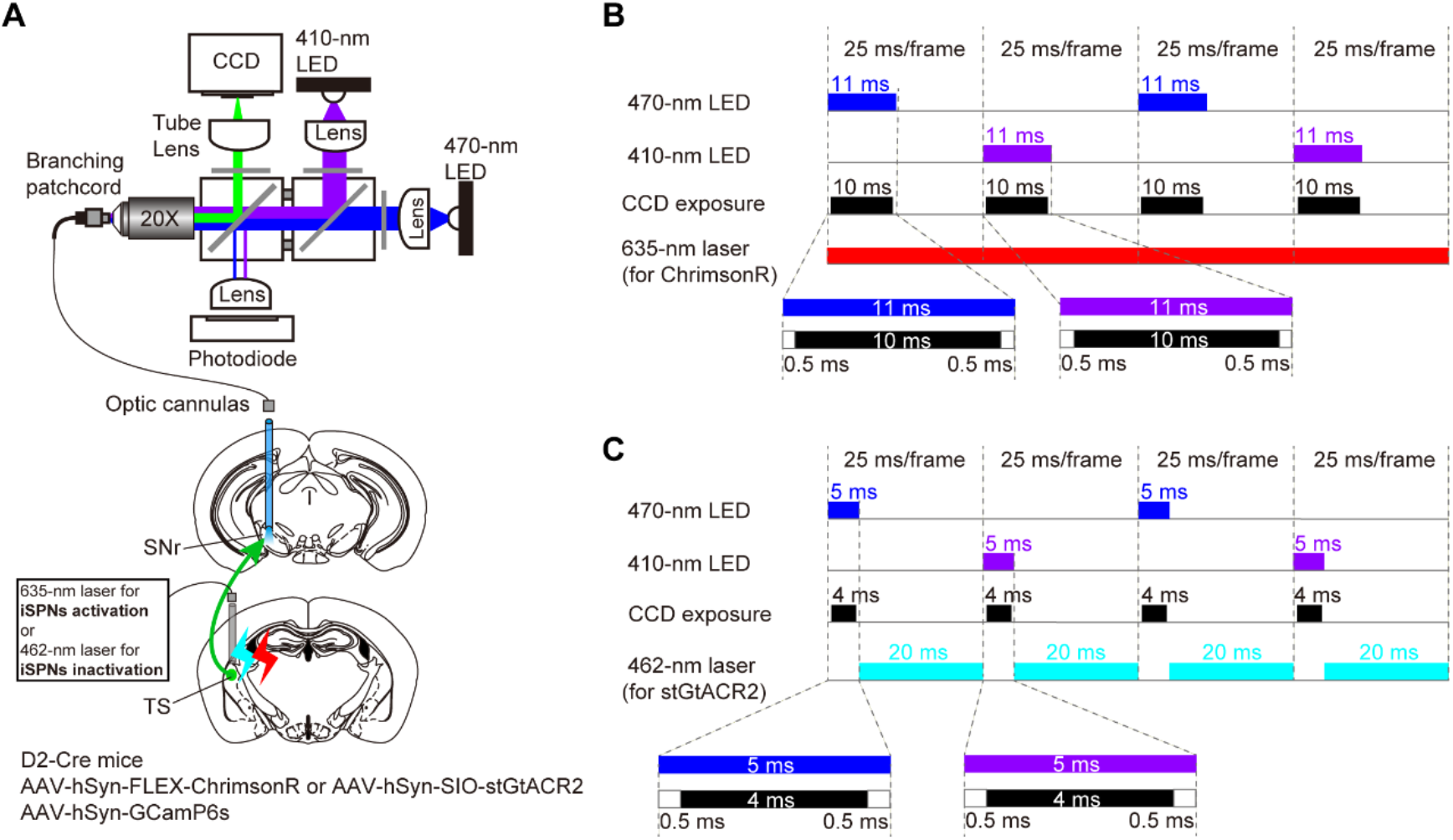
Time settings for fiber photometry recording and simultaneous optogenetic stimulation. (A) Schematic showing fiber photometry system and simultaneous optogenetic stimulation (see Methods). (B) Time settings for fluorescence excitation light (470 and 410 nm) and camera exposure for regular fiber photometry recordings, as well as for simultaneous optogenetic stimulation using 635 nm red laser for ChrimsonR-based activation. (C) Time settings for fluorescence excitation light (470 and 410 nm) and camera exposure during fiber photometry with simultaneous optogenetic stimulation using 462 nm blue laser for stGtACR2-based inactivation.

**Supplementary Figure 12.**
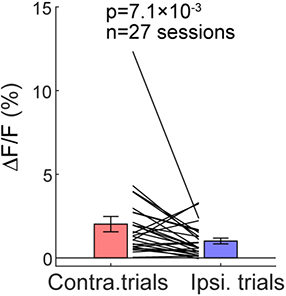
Axonal activity of dSPNs in SNr shows dominant preference for contralateral choices. Comparison of axonal activity between contralateral and ipsilateral correct trials without photostimulation. Each pair represents one recording session (n = 27 sessions from 12 mice, paired two-sided Wilcoxon signed rank test).

**Supplementary Table 1.**
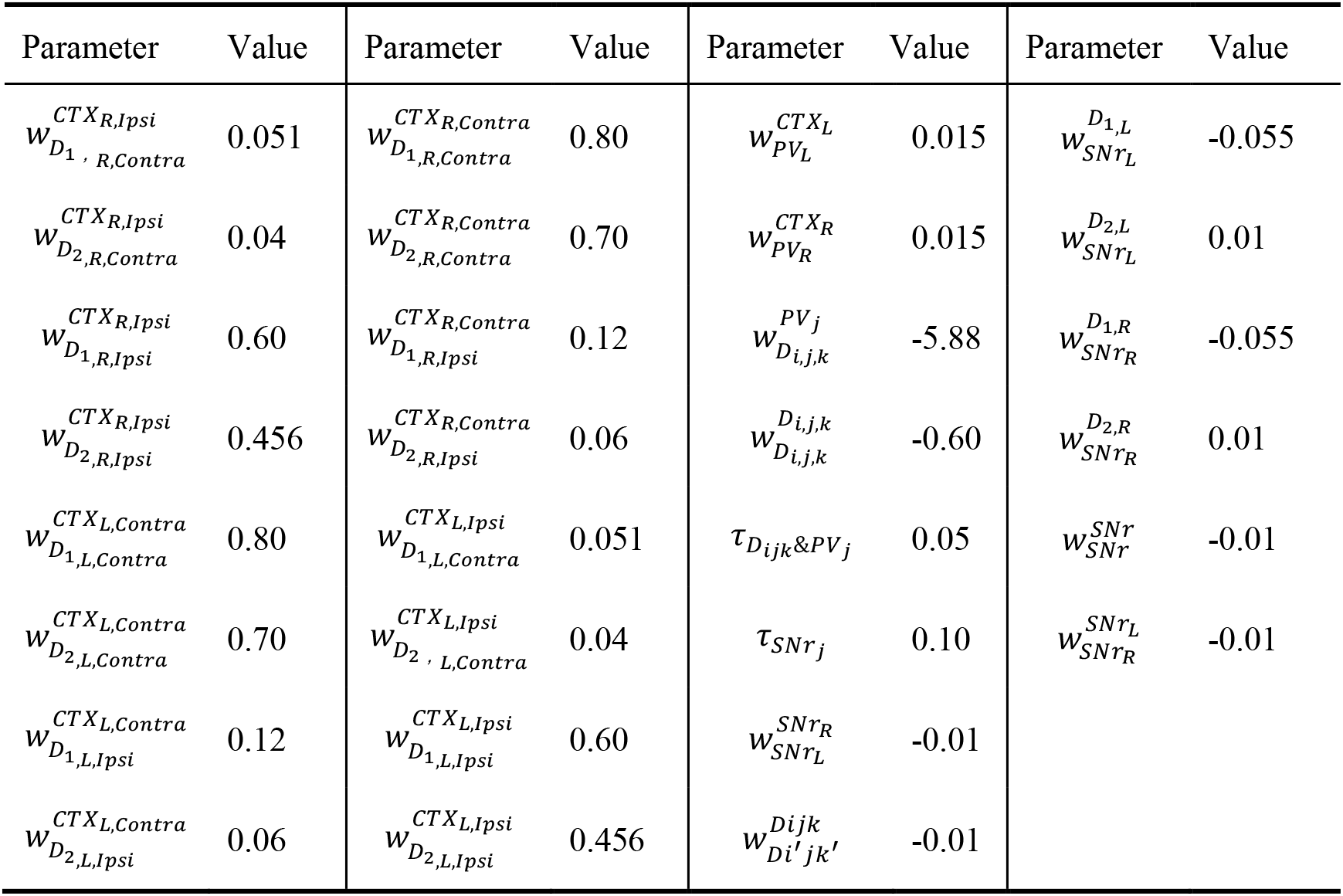
Parameters for the network model.

**Supplementary Table 2.** Optogenetic perturbation current of left hemispace (μA).

| Type | D1 activation | D2 activation | D1 inactivation | D2 inactivation | PV inactivation |
| --- | --- | --- | --- | --- | --- |
| $I^{stim}$ | 10.5±2.5 | 10.5±2.5 | -2.5±1.0 | -2.5±1.0 | -2.5±1.0 |

**Supplementary Video 1: Example video of optogenetic stimulation during behavioral task.** Video clip showing optogenetic activation of dSPNs in the right TS during the mouse performing the behavioral task. The OLED board on the right showing trial information and performance outcome. Note that the photostimulation in the third trials shown was accompanied by a wrong choice of the animal.

